# SEEKER: A genome-scale library-on-library screening platform for deciphering T cell recognition of antigen

**DOI:** 10.64898/2026.09.25.753432

**Authors:** Sicheng Li, Li Lin, Wenxu Wu, Binwen Liu, Huihui Ji, Zongyu Guo, Yaqiong Liu, Juncheng Wu, Bo Liu, Lisang Zhong, Xiaowo Wang, Huji Xu, Hai Qi

**Author notes:** Correspondence should be addressed to Hai Qi, Huji Xu or Xiaowo Wang.

## Abstract

Decoding which antigens activate a given T cell receptor (TCR) is a central challenge in immunology. We present SEEKER, a functional genome-scale library-on-library screening platform built on three elements: a Jurkat-derived T-APC cell (TAPCell) that combinatorially expresses one TCR and one peptide–MHC drawn from separate libraries together with an NFAT activation reporter; encapsulation of single TAPCells in a thermos-reversible hydrogel for isolated clonal expansion, so recognition is read out as intraclonal activation; and dual-asymmetric PCR that links TCR- and pMHC-encoding sequences into a heritable unit, enabling progressive hit enrichment over screening rounds. SEEKER interrogates ∼10^8^ TCR-pMHC combinations per run without prior knowledge. Screening joint-infiltrating CD8⁺ T cells from HLA-B*27⁺ ankylosing spondylitis patients against proteome-wide libraries validated 53 TCR– peptide pairs, uncovered extensive cross-recognition of self and common viral epitopes, and revealed highly polyreactive T cells recognizing up to 80 autoantigens, implicating virus-triggered poly-autoreactivity in multi-organ autoimmune diseases.

## Introduction

T lymphocytes recognize peptide antigens presented by major histocompatibility complex (MHC) molecules through clonotypic T cell receptors (TCRs). TCR-pMHC recognition underlies protective immunity to infection and cancer, but also drives autoimmune pathology when self-tolerance is broken. Identifying which antigens activate disease-relevant T cells is therefore critical for understanding immune pathogenesis and for rational design of antigen-specific therapies. With an estimated 10^12^ unique TCR clonotypes per individual^1,2^ and minimal clonal overlap between individuals, deciphering TCR-pMHC specificity remains a daunting technological challenge^3^.

Machine learning-based prediction of TCR-pMHC interactions is a promising approach but currently suffers from poor generalizability, owing largely to the scarcity of unbiasedly sampled, experimentally validated training data^4,5^. The inherent poly-reactive nature of TCRs, with each being able to recognize structurally distinct pMHC ligands through degenerate binding geometry, further confounds sequence-based prediction. Experimental data generation at scale is therefore a prerequisite for progress.

Several experimental strategies have been developed to map TCR antigen specificity. Conventional functional assays such as ELISPOT and intracellular cytokine staining^6^ require prior knowledge of candidate antigens and cannot be applied to unbiased discovery. DNA-barcoded pMHC multimers extend the throughput of antigen screening to approximately 10^3^ candidates in bulk populations^7,8^ or hundreds of antigens with single-cell receptor-antigen pairing^9,10^, but are constrained by the labor of individually assembling each barcoded multimer, the challenge of deconvolving interactions in large pools, and the limited set of MHC alleles amenable to recombinant folding.

Protein display libraries overcome some of those constraints. Baculoviral display of pMHC on insect cell surfaces enables screening of ∼10^5^ candidate antigens against purified, recombinant TCRs^11,12^. Yeast surface display of randomly synthesized peptide-MHC libraries at ∼10^8^ diversity, combined with deep sequencing of clones enriched over multiple rounds of selection, facilitates unbiased definition of TCR recognition landscapes and computational inference of natural ligands^13,14^. However, these display-based approaches select on binding affinity rather than functional activation, whereas high-affinity binders do not necessarily elicit TCR activation^15,16^. They require recombinantly produced, soluble TCR proteins formatted as multimers and, in the case of yeast display, allele-specific engineering of the MHC protein for stable surface expression, and screen only a single TCR per experiment.

Mammalian cell-based platforms address the functional gap by coupling antigen presentation to direct readouts of T cell activation. Several approaches convert pMHC complexes into the ligand-binding domain of synthetic signaling receptors, or exploit membrane transfer during T cell killing, in order to identify the pMHC targets of a given TCR^17–20^. These methods recover functionally validated interactions, but are limited to antigen libraries of 10^3^-10^4^, require high library redundancy, and interrogate only one TCR at a time. T-Scan extends this cell-based logic to proteome scale by delivering antigen libraries lentivirally into engineered HEK293T target cells expressing defined HLA alleles, using a granzyme B reporter to enrich cells recognized by CD8^+^ T cells, and identifying antigens by next-generation sequencing^21,22^. TScan-II adapted the platform to CD4^+^ T cells^23^. Both platforms require that TCRs be defined in advance, and each screen is antigen-centric but provides no direct readout of paired TCR-pMHC identity. More recently, lentiviral surface display of pMHC complexes has enabled receptor-antigen pairing at the single-cell level. The RAPTR platform exploits antigen-specific viral infection of TCR-expressing cells to link receptor and antigen identities through sequencing of the integrated viral barcode^24^, and ENTER-seq adds a simultaneous single-cell transcriptomic readout to this framework, permitting correlation of antigen specificity with T cell state^25^. However, both platforms require each member of the pMHC library to be individually cloned and packaged as a separate viral preparation, a bottleneck that limits the throughput to tens to hundreds of pMHC constructs. Moreover, these interaction-based methods do not accommodate iterative enrichment of functional TCR-peptide pairs over multiple rounds of screening, a prerequisite for identifying TCR-ligand combination at an omics scale.

It is highly desirable to develop a platform that can simultaneously screen large, diverse TCR libraries against proteome-scale pMHC libraries in a single pooled experiment with functional T cell activation as the readout. Such technology is particularly needed in autoimmune disease research, where the causative antigens are often unknown, disease-relevant T cell clones are numerous, and the specificity landscape may include self-reactive and microbially cross-reactive receptors.

Here we present SEEKER (Single-clone Encapsulation for Exhaustive Knowledge of Epitope-Receptor specificity), a platform that encodes TCR-pMHC combinations within individual Jurkat cells, identifies functional TCR-pMHC pairs based on cell activation due to intraclonal TCR-pMHC interactions, and achieves a combinatorial throughput exceeding 10^8^. We demonstrate the utility of SEEKER by profiling TCR poly-reactivities and screening joint tissue-infiltrating CD8^+^ T cells from ankylosing spondylitis patients against genome-wide autoantigen libraries.

## Results

### Rational design of TAPCell and SEEKER

An ideal TCR-pMHC screening platform should function as an apparatus that can take peptide epitopes, MHC alleles, and TCRs as three independent input variables to solve the equation of T cell activation in a high-throughput fashion. While sequencing is by far the highest-throughput readout, the central obstacle to varying those three variables independently in a sequencing-based platform is the fact that T cells are normally activated by antigen-presenting cells, and TCR and pMHC have to be genetically encoded in separate cellular/viral entities. Inspired by the observation that activated human T cells express both class I and class II MHC molecules and can therefore themselves function as antigen-presenting cells^26–28^, we considered the possibility of expressing exogenous TCR and pMHC in the same T cells to enable T cell auto-presentation and -activation in clonal culture (Figure 1A). For this, the pMHC component needs to be encoded in such a way that TCR and pMHC libraries could be varied independently and read out by sequencing. Two single-chain formats are available for this purpose. The single-chain trimer (SCT) fuses the peptide, β2-microglobulin (β2m), and MHC heavy chain into one polypeptide^29^. The single-chain dimer (SCD) links only the peptide and β2m, while the heavy chain is expressed as a separate entity^30^. Both formats allow the peptide library to be encoded in a single cassette together with a given MHC. The SCD is encoded in two shorter expression cassettes and additionally permits independent variation of the MHC allele identity and peptide pool (Figure 1B).

**Figure 1.**
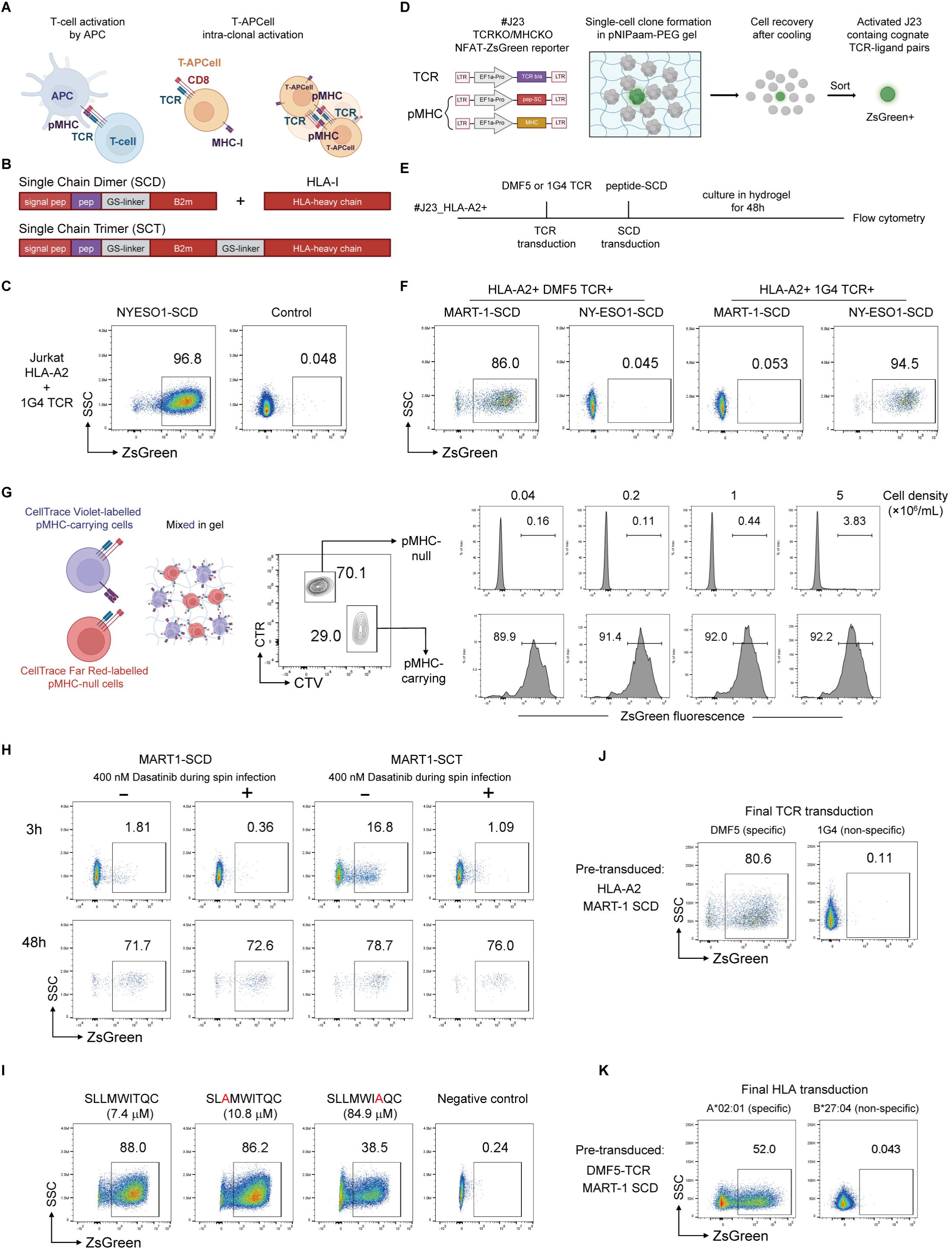
Development of the TAPCell system and SEEKER platform. (A) Schematic of conventional APC-mediated T cell activation (left) and intra-clonal auto-activation of T-APCells (right), which are Jurkat cells engineered to express an exogenous TCR and pMHC. (B) Schematic of the single-chain pMHC formats. The single-chain dimer (SCD) fuses the peptide to β2-microglobulin (β2m) through a flexible GS-linker and assembles with a separately expressed MHC heavy chain; the single-chain trimer (SCT) fuses peptide, β2m, and the MHC heavy chain into a single polypeptide. (C) Activation of Jurkat cells expressing the 1G4 TCR together with NY-ESO-1 SCD-HLA-A*02:01 reported by the NFAT-ZsGreen fluorescence. Control: Jurkat cells transduced with control vectors. (D) Schematic of the SEEKER workflow. Jurkat clone #23 is transduced with lentiviral vectors encoding the TCR β/α chains, a peptide-SCD library, and the MHC heavy chain. Single cells are encapsulated in thermo-reversible pNIPAAm-PEG hydrogel and allowed to expand as physically isolated clones. Cooling below the LCST liquefies the gel to release single cells, which are then sorted according ZsGreen expression. (E) Sequential transduction scheme for TAPCell validation. (F) ZsGreen profile of HLA-A*02^+^ J23 cells that expressed DMF5 or 1G4 TCR in combination with MART-1 or NY-ESO-1 SCD and were cultured as single-cell clones in gel. (G) Density dependence of activation of TAPCells (HLA-A*02^+^; 1G4 TCR; NY-ESO-1-SCD). Left: schematic of cell labeling and mixed culture in gel; Middle: the cytometry profile of cells after culture in gel; Right, percentage of activation reported by ZsGreen fluorescence at indicated total cell density encapsulated in gel. (H) Background reduction by Dasatinib treatment during spin infection (TAPCell: HLA-A*02^+^; DMF5 TCR; MART-1-SCD or -SCT). Dasatinib was washed out after infection. (I) TAPCell activation by affinity variant ligands (TAPCell: HLA-A*02^+^; 1G4 TCR; SCD of indicated peptide variants). Negative control contained irrelevant SCD. (J) Activation of TAPCells pre-transduced with indicated HLA-A*02 and MART-1 SCD to interrogate TCRs. In this example was specific DMF5 TCR versus mismatched TCR 1G4. (K) Activation of TAPCells pre-transduced with indicated DMF5 TCR and MART-1 SCD to interrogate HLA alleles. In this example was specific HLA-A*02:01 versus mismatched HLA-B*27:04.

To implement this design, we used CRISPR/Cas9 technology to disrupt the *Trac* and *Trbc* loci and genes coding HLA-A, -B, and -C in Jurkat cells that were also made to stably express an NFAT-ZsGreen-PGK-hCD8 (Figure S1A-C). The inclusion of hCD8 co-receptor increases the sensitivity of exogenously expressed TCR to its pMHC ligand^31,32^. The resultant master Jurkat clone, J23, was used for all subsequent experiments. We also disrupted HLA-A/B/C loci in 293T cells that were used throughout this study to package lentiviral vectors for subsequent transduction of J23 cells, preventing endogenous HLA in 293T cells from being carried by viral particles (Figure S1B).

J23 cells were then sequentially transduced with the 1G4 TCR, which recognizes NYESO-1 peptide in the context of HLA-A*02, and then with NYESO-1 SCD, and >96% of these cells became activated as judged by NFAT-ZsGreen fluorescence (Figure 1C). Similar results were obtained when 1G4 TCR and NYESO-1 SCT were combined (not shown). These data suggest that expression of exogenous TCR and pMHC on the same T cells is a viable strategy for TCR ligand identification. We term J23 cells that co-express TCR and pMHC in the SCD or SCT format the T-APC cell (TAPCell).

To be used for screening experiments, TAPCells would express random combinations of TCR and pMHC. In suspension culture, because TAPCells inevitably contacted neighboring cells and TCR on one TAPCell might be activated by pMHC on another TAPCell, it became difficult to ascertain the cognate correspondence between TCR and pMHC in ZsGreen^+^ TAPCells. To overcome this problem, drawing inspiration from agar-based semi-solid media used to isolate microbial colonies at high density, we searched for biocompatible hydrogel that could analogously encapsulate and physically isolate single mammalian cells from one another. Thermo-responsive poly(N-isopropylacrylamide)-poly(ethylene glycol) (pNIPAAm-PEG) hydrogels undergo reversible solution-gel transition at different temperatures and are biocompatible to permit clonal expansion of cells in isolation^33,34^. More specifically, the hydrogel remains in the liquid phase at 4°C, permitting uniform cell suspension, and solidifies at 37°C to physically separate each founder cell and its progenies as a colony from other founders. At the end of cultivation, the cell-containing gel can be liquified by cooling to release single cells for downstream analysis (Figure 1D). As shown in Figures 1E-F, TAPCells that contained the right TCR-SCD combination (i.e. DMF5 and MART-1, 1G4 and NY-ESO1) were activated in the gel with a ∼90% efficiency, while mismatched TCR-SCD combinations produced essentially no ZsGreen^+^ cells. Moreover, when we titrated TAPCells expressing the right TCR-SCD combination (1G4 and NY-ESO1) into J23 cells that only expressed 1G4 TCR, the activation efficiency remained very high when the cell frequency was as low as 0.01% (Figure S1D). With the TAPCell (1G4/NY-ESO1) frequency kept at 50% constant, when the total cell concentration in liquid gel was not higher than 5 million per ml, the bystander activation of J23 cells expressing only 1G4 TCR was less than 5% (Figure 1G). These events presumably resulted from random doublet formation and inter-clonal activation. Therefore, 5×10^6^ per ml gel was used as the upper limit of cell density for subsequent experiments.

The final step of assembling TAPCells involved spin infection with lentiviral vectors, and viral envelops may carry SCT or SCD proteins from 293T virus-packaging cells^24,25^, potentially causing unintended activation events during the spin infection. We thus tested whether Dasatinib, a clinically approved Src kinase inhibitor^35,36^, could be used during the 2-h spin infection to reduce such unintended activation. As shown in Figure 1H, Dasatinib treatment essentially eliminated the background activation events, which could be detected at 3 h, while it did not affect the efficiency of intended TAPCell activation read out 48 h after culture in gel.

To probe the sensitivity of our TAPCell platform to low-affinity peptide epitopes, we compared wildtype NY-ESO1 (SLLMWITQC) and its low-affinity mutant (SLLMWI<u>A</u>QC). The pMHC complex containing this mutant peptide exhibits an affinity to 1G4 TCR at approximately ∼85 μM kD, 10-fold lower than its wildtype due to the T→A substitution around the critical CDR3β contact site^25^. As shown in Figure 1I, approximately 88% and 38% TAPCells that expressed the wildtype and mutant SCD became activated, respectively, while the background was essentially null. Therefore, the TAPCell platform is sufficiently sensitive to detect ligands with very low affinities.

Finally, the modular design of this method conveniently provided the flexibility to either keep the pMHC as a constant while probing TCRs or keep the TCR-peptide combination as a constant while probing MHC alleles (Figures 1J-K).

Taken together, these data demonstrate that the hydrogel encapsulation combined with same-cell expression of TCR and SCD is highly specific and very sensitive to detect rare TAPCells that contain matching TCR-ligand pairs. We call this new method <u>S</u>ingle-clone <u>E</u>ncapsulation for <u>E</u>xhaustive <u>K</u>nowledge of <u>E</u>pitope-<u>R</u>eceptor Specificity or SEEKER.

### Fingerprinting a flu-specific TCR with its epitope mutation library

To understand TCR-ligand interactions and identify alternative ligands for a given TCR, library screening of related peptides is often required. To explore SEEKER capability of supporting large-scale library screening, we investigated the HLA-A*02:01-restricted epitope of influenza M1 protein, GILGFVFTL, in conjunction with its specific TCR (Figure 2A). Flu-M1 TCR and HLA-A*02:01 heavy chain were stably transduced into J23 cells in conjunction with a SCD library of peptides that keep the P2 isoleucine and P9 leucine residues of the native epitope constant but are otherwise mutated to have a Hamming distance of no greater than 5 (See Methods for detail).

**Figure 2.**
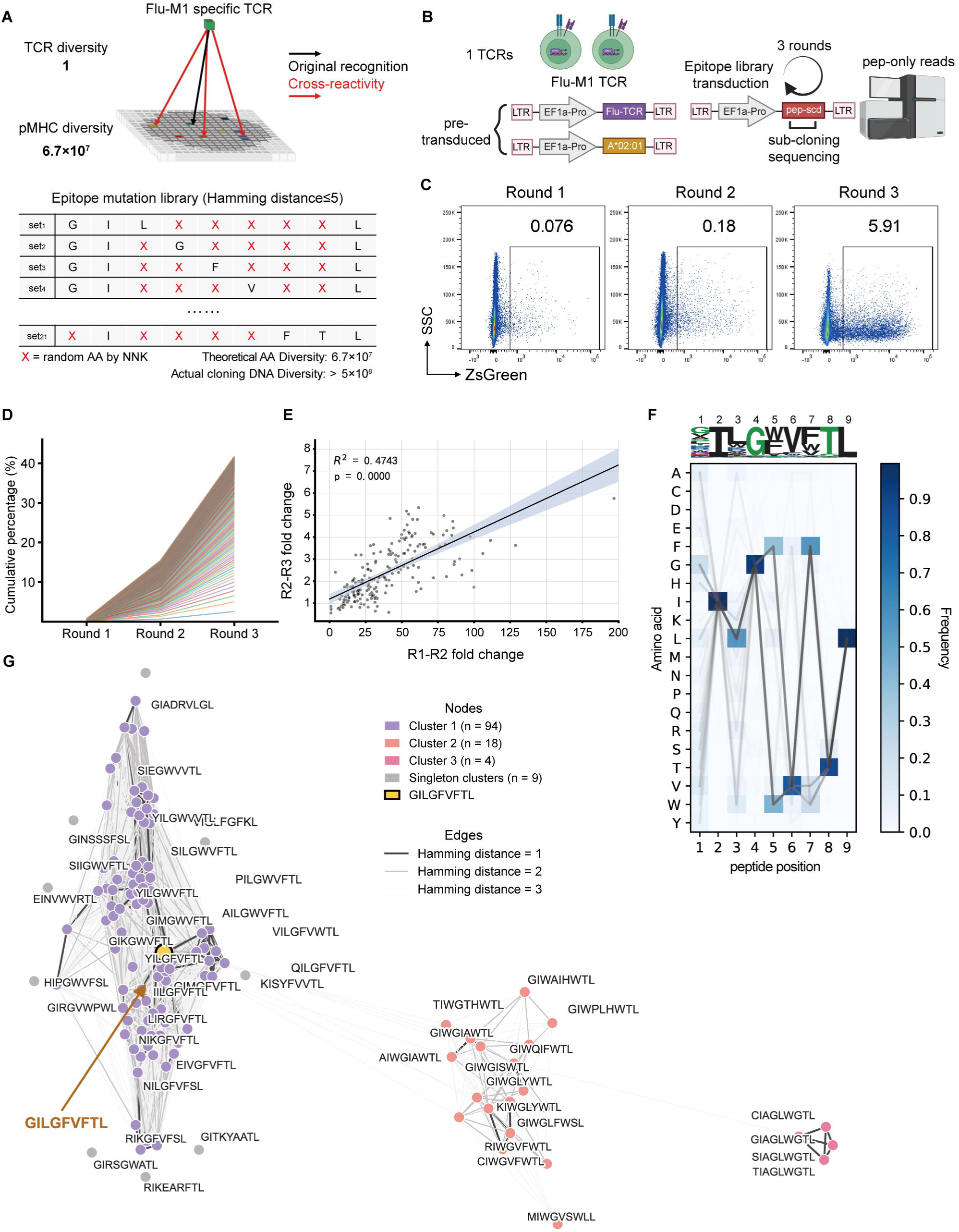
Epitope fingerprint of a flu-specific TCR by SEEKER. (A) Concept of recognition-fingerprinting (Top) and the design of an epitope mutation library (Bottom). (B) Schematic of the single-TCR SEEKER screen (TAPCell: HLA-A*02:01; Flu-M1 TCR; NNK-encoded SCD library as designed in (A)). (C) Cytometry profiles of ZsGreen^+^ cells at the end of each round. (D) Enrichment profile of top 200 sequences. (E) Correlation between R1–R2 and R2–R3 fold changes of top 200 sequences. (F) SeqLogo and amino acid usage matrix of top 126 peptides. (G) Sequence-similarity network of top 126 peptides. Edges connect peptides at the indicated Hamming distances.

To screen, the SCD library was transduced into 25 million J23 cells that uniformly expressed Flu-M1 TCR and HLA-A*02:01. ZsGreen^+^ TAPCells were recovered after 48 h, coding sequences of peptides were PCR amplified from the genomic DNA for making the next-round SCD library and for sequencing analysis (Figure 2B). After 3 rounds of screening, when the fraction of ZsGreen^+^ TAPCells surpassed 1% (Figure 2C), Peptide sequences were recovered. The pool of final top 200 sequences yielded 126 unique peptides after removal of non-productive and duplicated barcodes. These peptide sequences were progressively enriched across successive rounds of screening (Figure 2D). For any given sequence in this enriched final pool, its fold change between round 1 and 2 was proportionally correlated with its fold change between round 2 and 3 (Figure 2E, R²=0.4743). These results demonstrate stable and consistent enrichment expected of a robust screening process.

As illustrated in the SeqLogo plot and amino acid frequency matrix (Figure 2F), a Flu-M1 TCR recognition pattern emerged from those top 126 peptide sequences: minimal amino acid preference at P1, very strong preference for glycine (G) at P4, valine (V) at P6 and threonine (T) at P8, and marked preference for either phenylalanine (F) or tryptophan (W) at P5 and P7. When the 126 peptide sequences were clustered according to Hamming distance and sequence similarities, three major clusters were observed. One main cluster was prominent but could be further separated into several sub-clusters (Figure 2G, left). A smaller cluster sharing W at P3 had multiple different substitutions in other positions as compared to the main cluster (Figure 2G, middle), and a tiny cluster that shared a tripeptide motif of AGL at P3-5 (Figure 2G, right). These data are consistent with the fact that multiple residues within a peptide epitope are in contact with CDR3 in a cooperative manner^37^, and they demonstrate SEEKER is suitable for mapping TCR-ligand interactions.

### Probing TCR poly-reactivity with a fully randomized library

TCRs are inherently polyreactive^38,39^. We explored the utility of SEEKER to probe TCR poly-reactivity by testing DMF5 TCR, which recognizes MART-1-derived EAAGIGILTV peptide presented on HLA-A*02:01 and has a recognition landscape previously defined by a yeast-display strategy^40^. DMF5 was screened against a fully randomized 9-mer peptide library generated through NNK codon mutagenesis (Figure 3A). By the same protocol as in Figure 2B, three successive rounds of screening (initial input of 50 million cells) produced a recognition profile for the DMF5 TCR (Figure 3B-C). Classified according to the Hamming distance, two dominant clusters of 9-mer peptides were observed to contain either GXG (in blue) or DXG (in yellow) motif (Figure 3D). These clusters recapitulate GIG and DRG motifs at the same P4-P6 register previously reported^40^. SEEKER identified additionally smaller clusters containing uncharacterized motifs, including WFCCA and DLGXXV.

**Figure 3.**
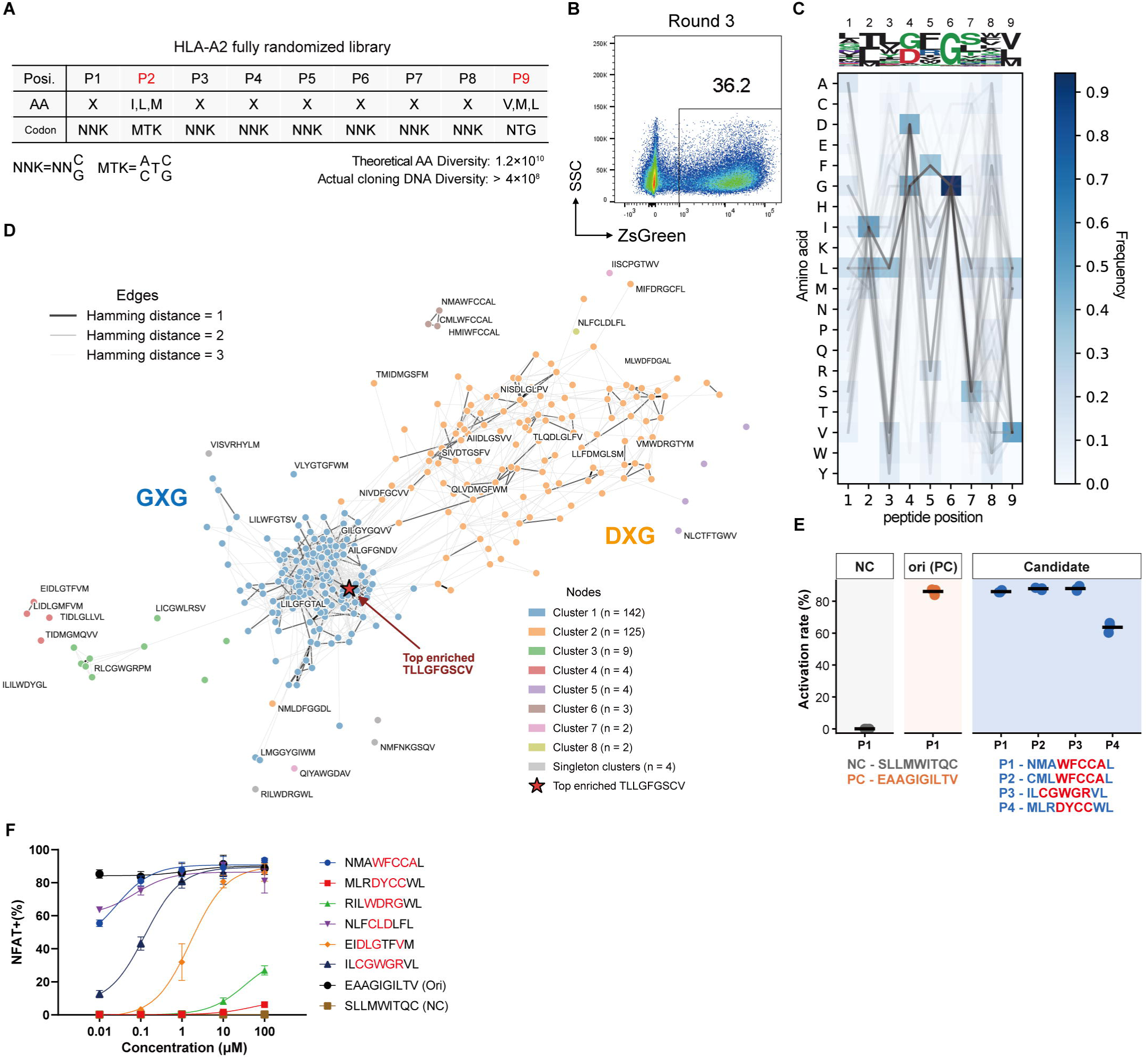
The polyreactive pattern of DMF5 TCR. (A) The design of a fully randomized HLA-A*02 9-mer peptide library. (B) Cytometry profiles of ZsGreen^+^ cells at the end of round 3. (C) SeqLogo and amino acid usage matrix of top-enriched 295 peptides. (D) Sequence-similarity network of top-enriched 295 peptides. Edges connect peptides at the indicated Hamming distance. (F) Dose responses of DMF5 TCR reacting to representative peptide ligands presented by K562 antigen presenting cells. Mean±s.e.m. is shown.

To validate these findings, representative peptides were expressed in their corresponding SCD format. The NMAWFCCAL peptide and its WFCCA analog activated TAPCells with a 87% efficiency as judged by the NFAT-ZsGreen^+^ frequency, the CGWGR-motif peptide achieved 89%, and the MLRDYCCWL peptide did 63% (Figure 3E). We further synthesized representative peptides and tested activation of DMF5 TCR-expressing J23 cells by HLA-A*02-expressing K562 cells pulsed with those peptides. As shown in Figure 3F, these peptides exhibited a range of potency spanning orders of magnitude: NMAWFCCAL was able to activate DMF5 at a nanomolar range, while MLRDYCCWL was able to activate DMF5 at 100 μM. Taken together, these data show that SEEKER is highly sensitive and can identify ligands of very low affinities, likely owing to its function-based readout of low background and high signal-to-noise ratio. These data also highlight the polyreactive nature of TCR, as one receptor can not only recognize related peptide ligands but also react to unrelated ligands of completely different sequences.

### Parallel fingerprinting of diverse TCRs by SEEKER

To test parallel screening of multiple TCRs in a single run, J23-A*02:01 cells were separately transduced with 4 distinct HLA-A*02-restricted TCRs (DMF5, 1G4, EBV, and Flu-M1). These cells were then mixed at equal ratios and subjected to screening against the same HLA-A*02-restricted NNK SCD library. Twenty-five million cells were allocated to cover each TCR in the first-round screen. During the 3^rd^-round of iterative screening, TCR-expressing J23-A*02 cells were color-coded with CTV and CTR and then transduced with SCDs enriched from the 2^nd^-round screen before being mixed for clonal culture in gel. The color code allowed us to purify activated TAPCells according to the TCR expressed (Figure S2A-B).

As illustrated in the SeqLogo plot and amino acid frequency matrix (Figures S2C), we recovered recognition motifs of all four TCRs in this mixed screening. The random 9-mer NNK-based SCD library has a theoretical peptide sequence diversity of ∼10^10^ (Figure 3A), while 25 million cells used to initiate the screen for each TCR could cover at most 0.25% of the entire sequence space. At this coverage, two independent screens are not expected to identify the same suite of peptides for a given TCR, but they should capture the same recognition fingerprint. Consistent with this expectation, very few peptides were identified in both screens for DMF5 (Figure S2D), while the mixed screen revealed the essentially same motif as the first screen (Figure S2E). Taken together, these data highlight the utility of NNK libraries for identifying TCR recognition motifs and demonstrate that the SEEKER platform can be used to fingerprint diverse TCRs in mixed screens.

### Linking TCR and pMHC into a single heritable cassette by in-droplet asymmetric PCR for high-throughput library-on-library screening

To identify rare TCR-pMHC pairs from omics-scale libraries, particularly when a large number of TCRs are involved, multi-round screening is required to progressively enrich cognate pairs from a pool of random combinations. Owing to the unique modular design of the TAPCell and hydrodel encapsulation system, SEEKER allows us a two-phase strategy to maximize effective throughput. In the first phase, a full-shuffle screen is conducted to search through all possible TCR-pMHC combinations, which require a throughput scaling as the multiplication product of TCR and pMHC libraries. In the second phase, two choices are available (Figure 4A).

**Figure 4.**
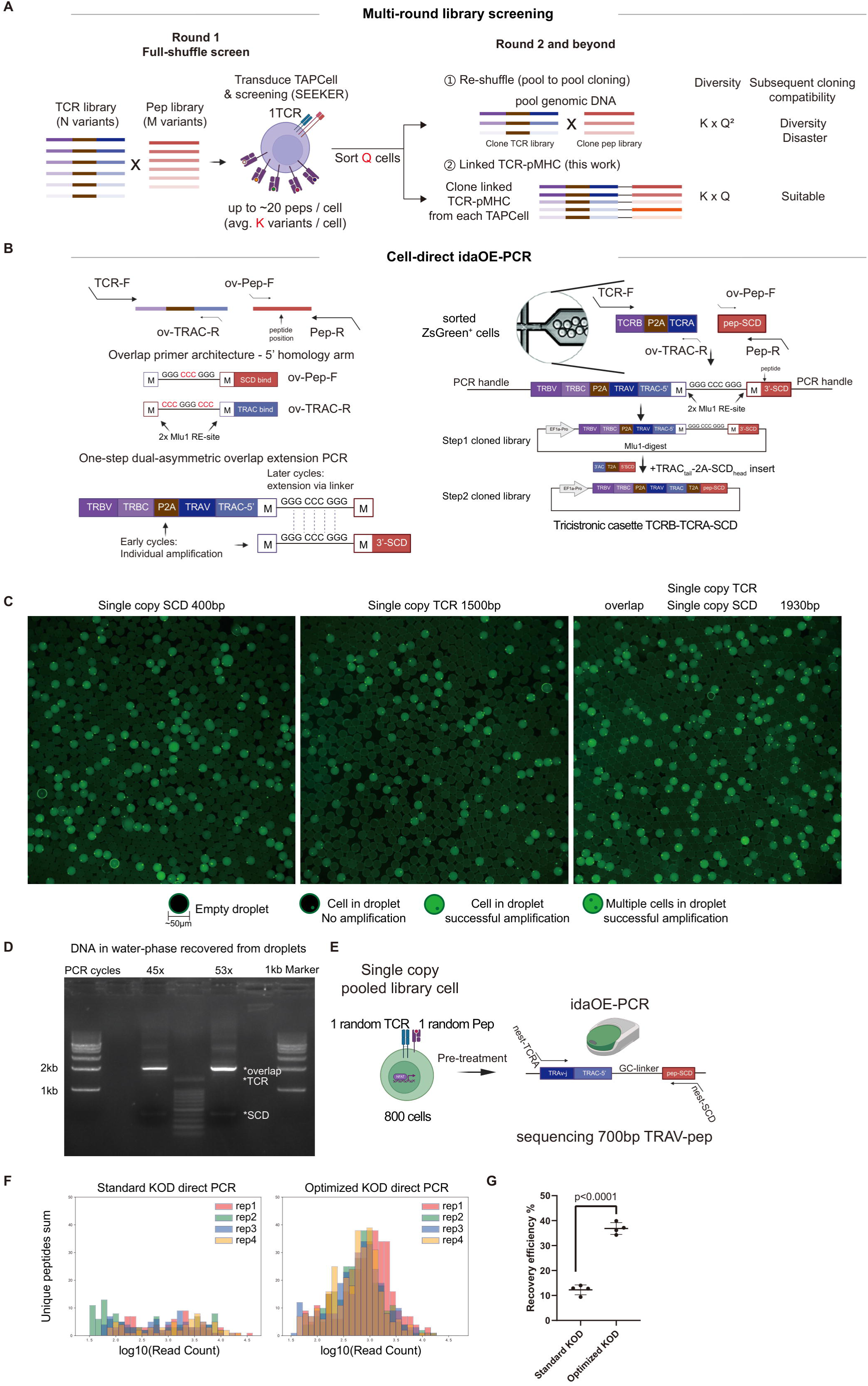
Heritable TCR-pMHC linked cassette for multi-round library-on-library screening. (A) Multi-round library screening strategies. (B) One-step and cell-direct idaOE-PCR for tri-cistronic TCR-SCD library construction. Left: primer positions to minimize the essential size of amplicons and principle of asymmetric overlap-extension PCR. Right: workflow schematic of tricistronic cassette construction. (C) Fluorescence images of droplets after amplification. Empty droplets, droplets with or without successful amplification are indicated. (D) An example of the linked TCR-SCD product amplified from DNA template in droplets. Overlap: linked product; TCR: intermediate amplification product of TCR; SCD: intermediate amplification product of the 3’SCD. (E) Workflow to test the efficiency of idaOE-PCR to recover TCR-SCD cassettes from 800 cells that each contained a single TCR and a single SCD. (F) Read-count distributions of unique peptides recovered by idaOE-PCR with standard KOD or optimized KOD conditions across four independent experiments. (G) Recovery efficiency under the two KOD conditions. Data presented as mean±s.e.m. *P*<0.0001 by two-tailed unpaired *t* test.

One is to pool NFAT-ZsGreen^+^ cells yielded from the first-round screen and clone all TCR and pMHC sequences as respective libraries to be used for subsequent full-shuffle screens just as the first-round. Once genomic DNA is pooled, the correspondence between a given TCR and its activating pMHC is lost, and any library re-cloned from the pool reverts to a random shuffle, which will lead to diversity disaster at the cloning step when the library size climbs (Figure 4A). The other possibility is to clone the TCR sequence and pMHC sequence from each TAPCell into one contiguous DNA molecule while the cell is still individually compartmentalized. The resultant library of linked TCR-pMHC cassettes can then be used as the substrate for subsequent screens (Figure 4A). This second strategy would enrich true cognate pairs faster and maximize the effective use of SEEKER throughput. We therefore implemented this latter strategy as a one-step, in-droplet dual-asymmetric overlap-extension PCR (idaOE-PCR) that amplifies the TCR and SCD cassettes from the genome of a single sorted NFAT-ZsGreen⁺ TAPCell and fuses them into a single amplicon (Figure 4B).

Two considerations shaped the design. First, the amplicon must be short enough to be generated efficiently from the single-TAPCell genome present in a droplet, yet must carry everything needed to rebuild a functional expression cassette. We searched TRAC for an internal position at which the constant region could be split, one that supports efficient priming, tolerates an appended homology arm, and lies as far 5’ as possible to minimize the amplicon length, while still leaving a 3’ segment that can be restored in trans during cloning (Figure 4B, top left). Meanwhile, the peptide-coding segment was likewise truncated to minimize amplicon size. At the site eventually chosen, only the 5’ TRAC portion of the TCR sequence travels through the droplet reaction, while the remaining TRAC-furin-T2A-SCD segment is supplied later from the vector. The resulting amplicon is ∼1.5 kb, among the longest reported for direct in-droplet amplification from single-cell genomic templates.

Second, fusion and subsequent reconstitution are accomplished at the same junction. We used an asymmetric, two-pair primer configuration^41^ (Figure 4B). One of the inner pair of overlap primers reads out of the TRAC break site, and the other reads into the 5’ end of the SCD cassette, while their 5’ homology arms consist of a high-efficiency GGGCCCGGG linker^42^ flanked by two MluI sites (Figure 4B, middle left). In the early PCR cycles, these primers generate the two half-products and, through the shared linker, template their fusion (Figure 4B, bottom left); a second pair of outer primers, present in large molar excess, then drives exponential amplification of the fused product, so that a single reaction yields the joined TCR-linker-SCD amplicon with no intervening purification or ligation. The linker is bracketed by MluI sites and serves as a temporary bridge. After the emulsion is broken and amplicons are recovered in bulk (Figure 4B, right), MluI excision removes the linker and generates the compatible ends into which the withheld TRAC-furin-T2A-SCD fragment is inserted. This restores an in-frame, tri-cistronic TCRβ-TCRα-SCD cassette (Figure 4B, right), allowing the linked library to re-enter SEEKER screening. Each cognate pair now segregates as a single genetic unit and is enriched across successive rounds rather than re-randomized at every round.

Realizing this reaction inside droplets required additional optimization. Genomic DNA within intact cells is poorly accessible, and although KOD polymerase tolerates crude templates, amplification occurred in only a fraction of cell-containing droplets, accompanied by non-specific products and droplet coalescence (Figure S3A). We found that supplementing the reaction with ET-SSB and additional dNTPs suppressed non-specific amplification and stabilized the emulsion. Heat-induced lysis followed by NotI digestion of genomic DNA further improved amplification efficiency from whole-cell templates and also prevented droplet fusion, likely by reducing the burden of intact genomic material. Together, these optimizations enabled successful amplification of genomic DNA in ∼90% TAPCells (Figure 4C). After PFO demulsification, the reaction products consisted mainly of the expected overlapping amplicons, indicating efficient linkage of the two loci (Figure 4D).

To evaluate performance, cells that each carries a single random TCR from a pool of 50 and a single SCD from a pool of 10^5^ random NNK peptides were generated by low-MOI transduction (Figure S3B-C). Out of the total diversity of 50×10^5^, we sorted 800 cells to PCR-link TCR and SCD (Figure 4E). A Hamming distance-based filter was devised to distinguish valid peptide barcodes from amplification and sequencing errors (Figure S3D-E). The optimized KOD protocol increased the recovery of input TCR-SCD combinations by more than 3 folds, and the recovered library showed good uniformity (Figure 4F-G).

### Identification of pMHC ligands for joint-infiltrating T cells of ankylosing spondylitis patients

Ankylosing spondylitis (AS) is a chronic inflammatory disease of the axial skeleton that progressively erodes spinal mobility, often leading to irreversible disability^43^. The disease exhibits the strongest HLA association known in human autoimmunity, as 90-95% of AS patients carry HLA-B*27, compared to <10% in unaffected populations^43,44^. It has been hypothesized that pathogenic CD8^+^ T cells recognize specific self-peptides presented by HLA-B*27, possibly triggered by cross-reactive microbial antigen^45^. However, identification of disease-driving or disease-associated autoantigen remains a daunting challenge. Yang and colleagues used HLA-B*27:05-restricted, yeast-displayed random peptide libraries to screen disease-associated TCRs, requiring antigen prediction based on peptide motifs identified^46^.

To SEEKER-screen disease-associated TCRs directly against the human proteome restricted by HLA-B*27, we first performed single-cell RNA sequencing (scRNA-seq) and TCR sequencing on cells recovered from the articular cartilage and periarticular tissues, which were obtained from AS patients undergoing hip-replacement surgery (Figure 5A). As expected, heavy infiltration of immune cells including T cells was seen across the joint, synovia, and the synovial fluid (Figure S4A-C). Within the T cell compartment, effector T cells, effector memory T cells, regulatory T cells, γδ T cells, and MAIT cells were readily identified (Figure 5B). Based on TCR clonal analysis, clonal expansion was detected predominantly in CD8+ T cells (Figure 5C; Figure S4D). Joint-infiltrating CD8+ T cells from our AS patients contained many more GZMK-expressing cells than GZMB-expressing cells (Figure 5D-E), consistent with the frequent predominance of GZMK expression in CD8+ T cells during chronic inflammatory diseases, in which GZMK activates complement^47,48^. Extensive clone sharing was observed across different sampling sites from the same patient (#2 and #3; Figure 5E). We curated the 100 most highly expanded CD8+ TCR clonotypes, comprising 49 from two patients with the HLA-B*27:04 genotype and 51 from two patients with the HLA-B*27:05 genotype (Figure 5F). Member cells of these clones predominantly expressed GZMK (Figure 5E).

**Figure 5.**
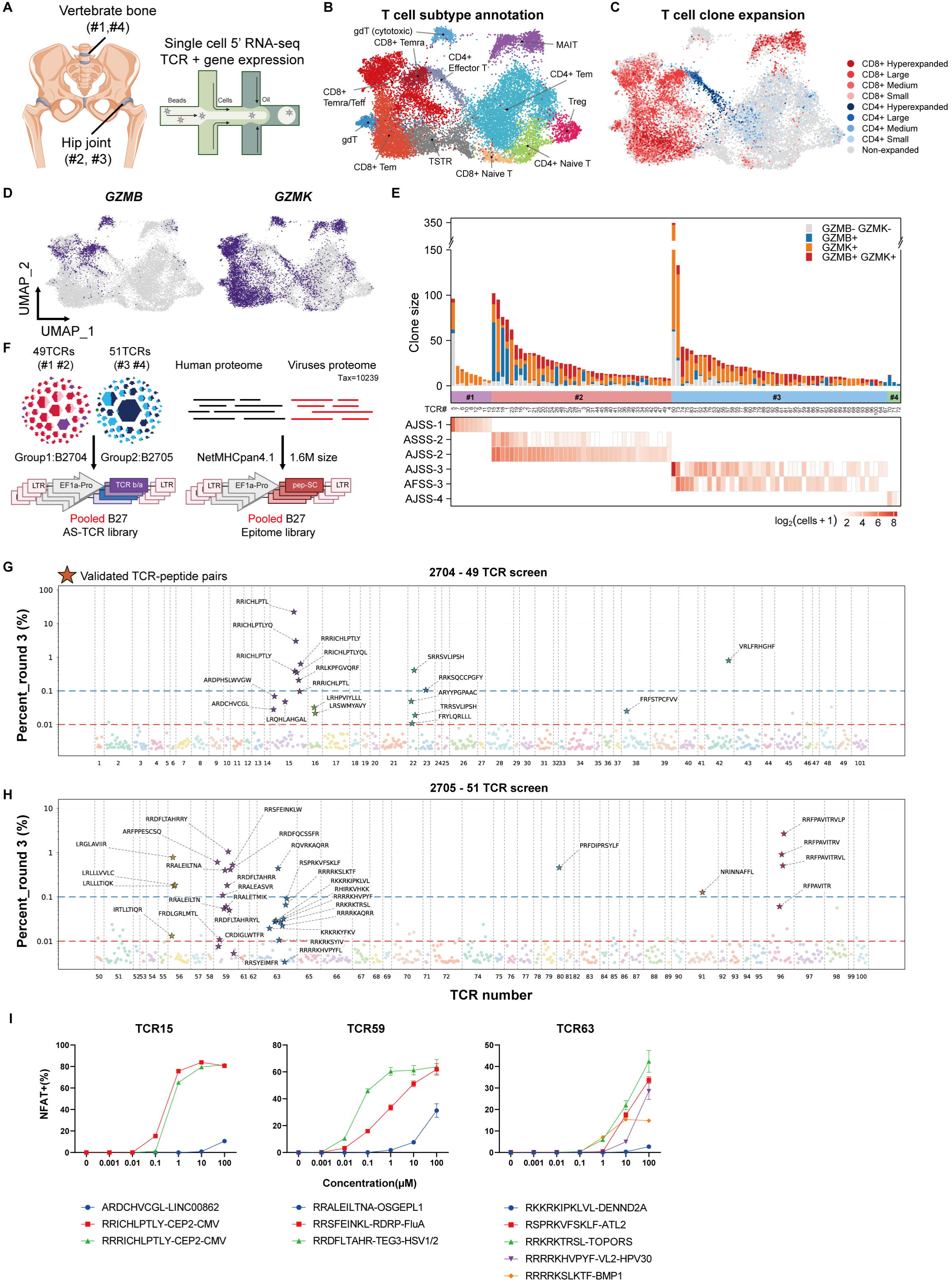
Ligand identification by SEEKER for disease tissue-infiltrating CD8^+^ T cells in ankylosing spondylitis. (A) Schematic of tissue sample sources and single-cell profiling to clone TCRs. (B) UMAP visualization of T-cell subtypes by scRNA-seq analysis. (C) Patterns of clonal expansion of T cells, with clones categorized according to expansion size. (D) Patterns of GZMB and GZMK expression in T cells. (E) Characteristics of the 100 most expanded CD8^+^ T cells selected for ligand identification, arranged according to sample source, clone size, GZMB and GZMK expression patterns. Top: clone-size distribution, color-coded by GZMB/GZMK expression; Bottom: per-sample clone distribution on a log2(cell count+1) scale. Sample source identified as AJSS, joint; ASSS, synovium; AFSS, synovial fluid. (F) Schematic of the library-on-library screen, showing that TCRs from two HLA-B*27:04 patients and two HLA-B*27:05 patients are respectively pooled as an AS-TCR library, and an SCD library of human and viral proteomes is built based on netMHCpan4.1 prediction. (G and H) Peptide-enrichment landscapes of the final round for the HLA-B*27:04 group (G) and the HLA-B*27:05 group (H). Each point represents a peptide hit for the indicated TCR. Validated peptides are star-labeled. (I) Dose responses of three poly-specific TCRs (TCR15, TCR59, and TCR63) to indicated peptides presented by HLA-B*27-expressing K562 cells. Data are plotted as mean±s.e.m. of triplicate.

Next, we screened these TCRs against their respective HLA-B*27:04-or HLA-B*27:05-restricted SCD library that covers proteomes of human and all known viruses (tax: 10239) in 8-, 9-, 10-, 11-, 12-mer formats and reaches a size of approximately 1.5×10^6^ entries (Figure 5F). Because each patient carried 6 class I HLA alleles, only 1 out of 6 TCRs was expected to recognize B*27:04- or B*27:05-presented ligands, assuming balanced HLA restriction without any pre-enrichment.

J23-HLA-B*27 cells were transduced with the TCR library at a low MOI to favor single-copy integration and balanced expression (transduction efficiency adjusted to ∼5%; Figure S4E). The resulting 6×10⁸ TCR-expressing cells were subsequently transduced with the SCD library at near-saturating efficiency (∼100%). After 3 rounds of screening to enrich specific sequences, PacBio sequencing of the libraries identified multiple pairs of TCRs and corresponding peptides being enriched (Figure 5G-H). To confirm the authentic TCR-peptide pairings, for each TCR, we synthesized the SCD forms of the top candidate peptides together with additional peptides showing sequence similarity, and tested their ability to stimulate J23 cells expressing the corresponding TCR. We found that 6 out of 49 HLA-B*27:04-restricted TCRs recognized 9 self-peptides and 10 viral peptides, while 6 out of 51 HLA-B*27:05-restricted TCRs targeted 16 self-peptides and 18 viral peptides (Figure S4F-G).

Among these 12 TCRs, 3 (TCR15, TCR59, TCR63) exhibited poly-specificity, recognizing 10 or more antigens. They cross-recognized both viral and self antigens, which contained epitopes showing some similarities (Figure S4H). We synthesized representative peptides for each of the three TCRs and pulsed them onto K562 cells expressing the corresponding HLA-B*27 alleles as APCs. These peptides readily activated J23 cells expressing the corresponding TCRs. Interestingly, viral peptides were generally more potent than self peptides. (Figure 5I). These observations suggest that, despite substantial differences in the full-length sequence, core motifs could make large contribution to TCR reactivity.

### Deep profiling of polyreactive TCRs that recognize both self and viral antigens

Among all TCRs tested, TCR59 and TCR63 exhibited the broadest cross-reactivity. To characterize their recognition landscapes in greater depth, we expressed TCR59 and TCR63 individually in J23-HLA-B*27:05 cells and screened them against the same proteome SCD library. Such single-TCR screens do not require assembly of the tricistronic cassette and consequently allow TAPCells to express higher levels of TCR and SCD and perform at a greater sensitivity.

After two rounds of enrichment, peptide-encoding inserts were recovered by cloning and identified by sequencing (Figure 6A). We then individually validated the top 192 candidate peptides for each TCR in J23 cells expressing the TCR at single-copy levels. For TCR59, 107 of the 192 peptides (55.7%) showed positive activation (>5%), 90 (46.9%) were strong activators (>30%). For TCR63, 132 peptides (68.8%) were positive, 131 (68.2%) were strong activators (Figure 6B). To assess the impact of TCR expression level, we repeated the validation of all 192 TCR59 candidates in J23 cells expressing a high copy number of TCR59. Activation was highly concordant between the two expression settings (Pearson r = 0.98; Figure S5A), with only two additional peptides becoming detectable under high-copy condition, indicating that the single-copy system captures nearly the full ligand repertoire.

**Figure 6.**
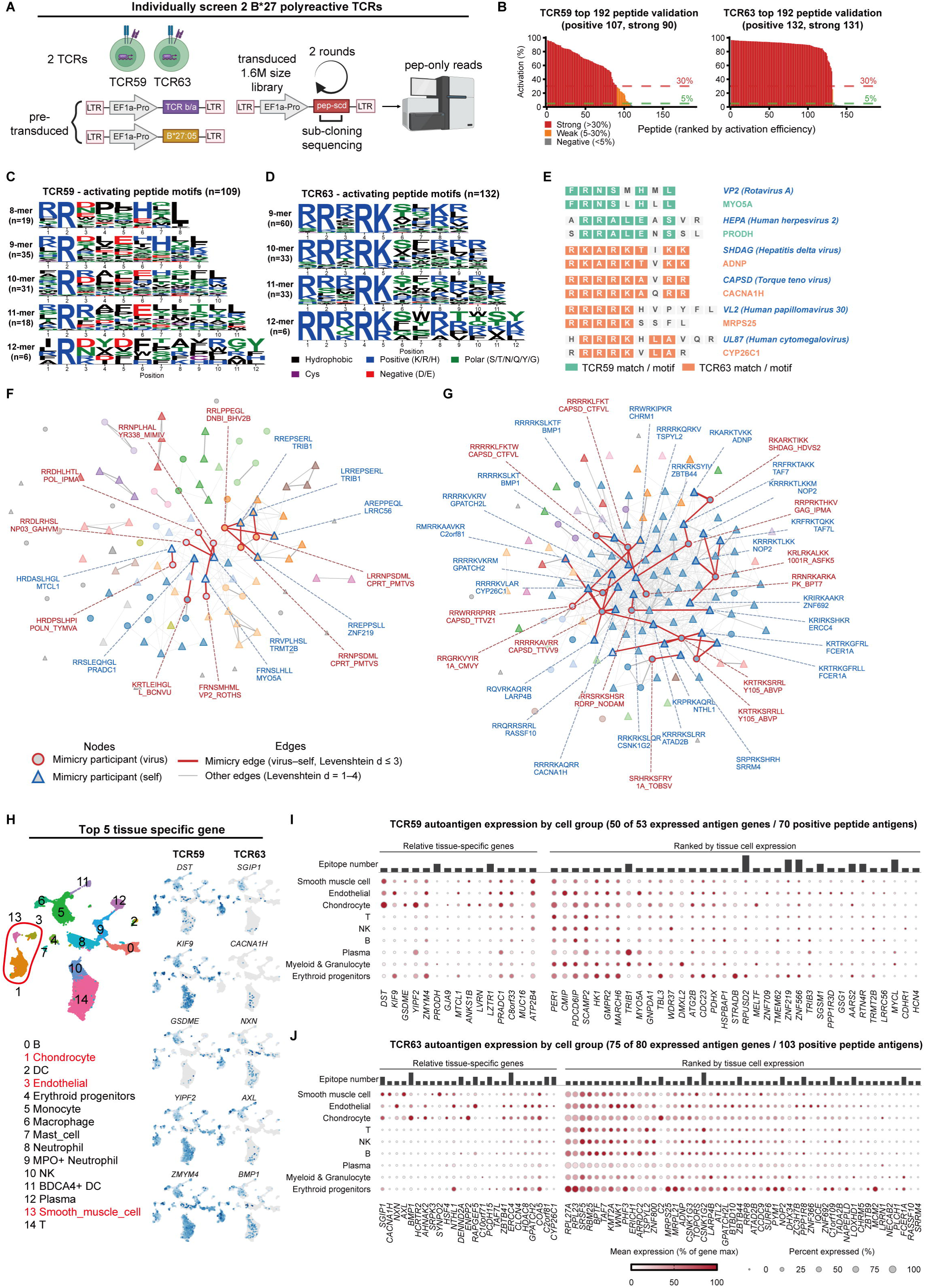
Deep profiling of highly polyreactive TCRs of disease tissue-infiltrating CD8^+^ T cells in ankylosing spondylitis. (A) Schematic of the focused screen for TCR59 and TCR63. (B) Individual validation of the top candidate peptides for each TCR in the NFAT reporter assay. Peptides (n = 192 per TCR) are ranked by activation efficiency. Bars are colored by response class—strong activators (>30%), weak activators (5–30%), and non-activators (<5%)—and dashed lines mark the 5% and 30% activation thresholds. (C and D) Motifs of activating peptides for TCR59 (C) and TCR63 (D), shown as sequence logos stratified by peptide length. Letter height is proportional to residue frequency, and residues are colored by physicochemical class. (E) Representative virus–self peptide pairs showing molecular mimicry. Viral peptides (viral protein and species) are aligned with homologous human self peptides (gene symbol); positions matching the TCR59 motif are highlighted in teal and those matching the TCR63 motif in orange. (F and G) Sequence-similarity networks of the mimicry-participating activator peptides for TCR59 (F) and TCR63 (G). Peptide sequences were hierarchically clustered, with nodes arranged and colored by cluster membership. Nodes represent peptides derived from viral (circles) or self (triangles) proteins, and red edges connect virus–self pairs within a Levenshtein distance of three. (H) UMAP visualization of the AS joint-tissue scRNA-seq data with annotated cell types, and feature plots of the top tissue-specific autoantigen genes for TCR59 and TCR63 on a grey-to-blue expression scale. (I and J) Autoantigen expression by cell group for TCR59 (I) and TCR63 (J). Dot color indicates mean expression relative to each gene’s maximum across cell groups, and dot size indicates the fraction of expressing cells; tissue-specific genes are grouped on the left, and the remaining genes are ranked by tissue-cell expression on the right, with the epitope count per gene shown above the panel.

Motif analysis of the confirmed peptides showed that TCR59 recognizes 8- to 12-mer peptides with diverse yet related sequence patterns. Apart from the P2 arginine that serves as the canonical HLA-B*27 anchor, no strong positional constraint was apparent, and the favored residues shifted across length classes. In contrast, the confirmed TCR63 peptides were restricted to 9- to 12-mers and carried, beyond the shared P2 anchor, a highly positively charged N-terminal stretch with an almost invariant Arg–Lys pair at positions 4 and 5 (Figure 6C-D).

The confirmed peptides derived from both self and viral antigens—70 self-derived and 39 viral peptides for TCR59, and 103 self-derived and 29 viral peptides for TCR63. For TCR59, mimicry was observed between rotavirus A VP2 and human MYO5A, and between human herpesvirus 2 HEPA and human PRODH. For TCR63, hepatitis delta virus SHDAG and human ADNP differed by a single residue, as did torque teno virus CAPSD and human CACNA1H, with additional mimicry observed between human papillomavirus 30 VL2 and MRPS25 and between cytomegalovirus UL87 and CYP26C1 (Figure 6E). To systematically survey such relationships, we constructed sequence-similarity networks for the confirmed peptides of each TCR, with self–viral peptide pairs of Levenshtein distance ≤3 highlighted in red, revealing multiple such pairs for both TCRs (Figure 6F-G). Extending this survey with relaxed similarity criteria showed that the top mimicry targets for each TCR—five human genes per TCR—were dominated by common human viruses, most prominently the herpesvirus family Orthoherpesviridae (Figure S5B). These observations highlight viral mimicry as a potential mechanism to activate highly polyreactive T cell clones such as TCR59 and TCR63, which have an increased potential to promote autoimmune pathology.

Autoimmune diseases including AS may progressively affect multiple organs, owing in part to activation of T cells reactive to universal autoantigen and/or multiple T cells reactive to organ-autoantigens from different organs. In this regard, highly polyreactive T cells can be particularly dangerous, because even a single clone may react to multiple universal and organ-specific autoantigens. We therefore characterized potential tissue and cell origins of autoantigens that TCR59 and TCR63 react to. First, confirmed TCR59 and TCR63 peptides were mapped to 53 and 80 autoantigens, respectively. To assess the relevance of these autoantigens, we examined their expression in our single-cell transcriptomic dataset of joint-associated tissues from AS patients (Figure 6H), in which clusters 1, 3, and 13 were allocated to tissue cells, including smooth muscle cells, endothelial cells, and chondrocytes. For TCR63, several top autoantigens, including SGIP1, CACNA1H, NXN, AXL and BMP1, were mainly expressed by these tissue-cell clusters, as were select TCR59 autoantigens such as DST, KIF9, and GSDME, while other TCR59 candidates displayed a broader expression pattern (Figure 6I-J).

In summary, by SEEKER screening our work identifies autoreactive CD8⁺ T cells in joint tissues of HLA-B*27-carrying AS patients, uncovers highly polyreactive TCRs that can recognize epitopes derived from both common viruses and self antigens, and raises the possibility that viral infection may activate polyreactive T cells that recognize multiple autoantigens, inducing or exacerbating pathology of multiple organs in autoimmune diseases.

## Discussion

The central obstacle to decoding TCR specificity at scale is the fact that a T cell is normally activated by a separate antigen-presenting cell, and the three variables that define recognition, peptide, MHC allele, and TCR, are encoded in two different cellular entities and cannot be freely varied in a sequencing-based screen. SEEKER collapses the two cells into one, co-encoding a TCR and a pMHC in a single Jurkat reporter cell. While co-expression in suspension allows a TCR on one cell to be engaged by pMHC on a neighbor, destroying the correspondence between receptor and ligand, single-clone encapsulation in thermo-reversible hydrogel resolves this issue by confining activation to a founder cell and its progeny. Encoding the antigen in a SCD rather than a SCT format keeps the peptide pool and the MHC allele independently variable, so the same library can be redirected to a different allele, a different species, or to class II MHC without redesign. Reading activation through an NFAT reporter rather than binding affinity selects for engagements that actually signal, and the near-zero background makes iterative enrichment possible. Finally, in-droplet linkage converts a functional hit from a transient property of a cell into a heritable genetic unit that can re-enter the screening process. The combination of these design elements has given SEEKER its unique ability to allow TCR and antigen libraries both be scaled at will.

The sensitivity of SEEKER is evident in the fingerprinting experiments. Screening a single TCR against fully randomized NNK libraries, SEEKER recovered two recognition classes previously defined for DMF5 by yeast display at the same P4–P6 register, and additionally returned motifs that had not been described but can be verified using synthetic peptides. Peptides spanning three orders of magnitude in potency were recovered from the same screen, including ligands at the low-affinity end.

The capacity for genuine, omics-scale library-on-library is a distinguishing feature of the SEEKER platform, and it rests on iterative enrichment. Co-culture screens using polyclonal T cells against antigen libraries do not directly return TCRs and peptides in matching pairs. Virus-display approaches such as RAPTR and ENTER-seq return pairs at single-cell resolution, but each member of the pMHC library must be individually cloned and packaged^24,25^, and, more importantly, there is no mechanism to convert the pairs identified in one round into a linked library for the next. Without that step, screening the combinatorial space of a genome-wide antigen library against a large TCR panel would become intractable. By the same token, the apparent throughput, the product of TCR number and antigen library size in a single round of screening, is not a good measure of what a platform can actually discover. As demonstrated here, even when we set SEEKER to operate at a relatively limited throughput of ∼10⁸, well below that of interaction-based methods, it returned a substantial body of TCR-ligand pairs from genome-wide libraries with no prior enrichment and no a priori knowledge, because each round narrows the search space for the next.

Applying SEEKER to ankylosing spondylitis illustrates what this makes possible. We screened 100 clonally expanded, joint-infiltrating CD8⁺ TCR clonotypes from four HLA-B*27⁺ patients against HLA-B*27:04- and B*27:05-restricted libraries, which spanned the human proteome together with the proteomes of all known viruses, in 8- to 12-mer formats. The screen returned 12 TCRs with validated ligands and 53 TCR– peptide pairs in aggregate, a volume of paired specificity data that, to our knowledge, have not been produced from single screening studies.

Our results provide direct evidence that many of the AS disease-associated TCRs in the joint tissues recognize self peptides. Interestingly, some of those receptors also recognize epitopes derived from common viruses carried in human population such as CMV. Although it is difficult, at this stage, to ascertain precisely the roles of those viral reactivities play in AS disease pathogenesis, it is plausible that they contribute to infection-associated flares and exacerbation of the disease. Furthermore, deep profiling by SEEKER reveals a subset of those receptors as extremely polyreactive, recognizing self epitopes derived from as many as 80 autoantigens, alongside viral epitopes sharing the same core motifs. Cells of this kind would have been very difficult to observe by any existing approach, but their presence among the most expanded clones in the inflamed tissue suggests that they may be common in autoimmune lesions rather than exceptional. Interestingly, these cells also express GZMK, a proinflammatory granzyme increasingly recognized as a key pathogenic factor in allergy and autoimmune disease^47,48^. Because a single polyreactive receptor can recognize epitopes derived from antigens expressed in unrelated tissues, T cells carrying such antigen receptors could in principle sustain inflammation at several anatomical sites, thus potentially explaining why autoimmune diseases so frequently begin from one tissue or organ and spread to involve multiple organs.

Because human proteome is of a finite size, the SEEKER platform is particularly useful for identifying peptide ligands of self-reactive CD8 T cells in autoimmunity. However, SEEKER is not limited to this single purpose but should be readily applicable to research of cancers and infectious diseases. Class II libraries can be screened with the same workflow in principle. That said, current SEEKER also has several limitations. First, current antigen libraries are assembled from computationally predicted HLA-binding peptides, leaving open the possibility that some ligands identified here may not be processed and presented under physiological conditions. Second, although in-droplet linkage removes the throughput ceiling limited by molecular cloning, the screen remains bounded by the number of cells that can be cultured. Third, post-translationally modified and non-genetically encoded antigens cannot be represented in a genetically encoded library.

In summary, SEEKER establishes a functional, prior-knowledge-free route to TCR-ligand discovery in which both receptor and antigen are screened as libraries, and its application to ankylosing spondylitis demonstrates that the approach can be brought directly to human diseases. We expect this platform to be broadly useful for studies of immune-mediated diseases, infections and cancers.

## Methods

### EXPERIMENTAL MODEL AND STUDY PARTICIPANT DETAILS

#### Study participants

The study was approved by the institutional review board, and all patients provided written informed consent in accordance with the Declaration of Helsinki before enrolment. The clinical protocol was reviewed and approved by the Clinical Research Ethics Committee of Shanghai Changzheng Hospital (Ethics Approval Number: 2022SL028). Synovial fluid and joint tissue samples were obtained from patients with ankylosing spondylitis undergoing total hip arthroplasty or spinal fracture internal fixation at Shanghai Changzheng Hospital. Written informed consent was obtained from all donors.

### METHOD DETAILS

#### Cell line construction

HEK293T cells were cultured in DMEM. To generate HLA-I knockout (KO) 293T clones, HEK293T cells were transfected with the pL-CRISPR.EFS.GFP vector carrying sgRNAs targeting HLA-A/B/C loci. After transfection, single cells were sorted, and clones with complete HLA-I KO were verified by flow cytometry.

To generate Jurkat cells deficient in TCR and HLA-I, 1–5×10⁶ cells were electroporated with the pL-CRISPR.EFS.GFP plasmid containing specific sgRNAs. TCR KO was verified by both flow cytometry and RT-PCR/sequencing, while HLA KO was validated by flow cytometry. A lentivirus carrying the NFAT-zsGreen-pgk-hCD8 cassette was used to transduce Jurkat cells, and a master clone, J23, was selected for its superior sensitivity to activation by exogenous TCR. J23 clone is then transduced to stably express HLA-A*02 or HLA-B*27 as appropriate for subsequent experiments and referred to as J23-A*02 or J23-B*27, respectively.

Sequences of guide RNAs used are as follow:

HLA-A-sgRNA-1: CGGCTACTACAACCAGAGCG HLA-B-sgRNA-1: AGATCACACTGACCTGGCAG HLA-C-sgRNA-1: AGGTCAGTGTGATCTCCGCA TCRa-sgRNA-1: AGAGTCTCTCAGCTGGTACA TCRa-sgRNA-2: TGTGCTAGACATGAGGTCTA TCRb-sgRNA-1: GCAGTATCTGGAGTCATTGA TCRb-sgRNA-2: GGAGAATGACGAGTGGACCC

#### Lentivirus production

For lentivirus production, HLA-I KO HEK293T cells in each 6-well plate were transfected with 1.25 μg viral expression vector, 0.3125 μg pMD2.G, and 0.9375 μg psPAX2 using Mirus-IT2 (Mirus Bio). The medium was replaced after 24 h, and viral supernatant was collected at 48 h. The virus was filtered through a 0.45 μm PES filter (Millipore), concentrated with 4×Lenti-X Concentrator (Takara) following the manufacturer’s protocol, and stored at -80°C as 30×concentrated stock. For large-scale lentivirus production, all transfection components were scaled up proportionally at identical ratios.

#### Lentiviral transduction

For routine lentiviral transduction during cell line construction, 1×10⁵ Jurkat or K562 cells were incubated with 1 mL unconcentrated lentivirus for 24 h. For certain experiments to achieve single-copy integration in T cells transduced, viral preparations were serially diluted to achieve a low transduction rate (<15% positive). For SEEKER (single-clone encapsulation for exhaustive knowledge of epitope– receptor) screening experiments, concentrated virus was added to complete RPMI-1640 supplemented with 400 nM dasatinib. Cells were seeded at 2×10⁶ cells/mL, centrifuged at 1,500 ×g and 32°C for 2 h, and washed twice with complete medium after transduction to remove residual virus particles and dasatinib.

#### 3D cell encapsulation culture in PNIPAAm-PEG hydrogel

PNIPAAm-PEG hydrogel (MebioGel) was dissolved in complete medium to a final concentration of 10% (w/v) following the manufacturer’s protocol. Jurkat cells were counted and resuspended in complete medium and mixed with chilled hydrogel solution at a 1:10 ratio on ice. The cell-gel mixture was then injected into pre-warmed complete medium and allowed to solidify by temperature-induced gelation. Post-culture, hydrogel culture flasks were placed on ice to liquefy the hydrogel. Released cells were washed twice with complete medium before flow cytometry analysis.

#### NNK peptide-SCD library design and construction

Primers, oligonucleotide pools, and gene fragments were ordered from GenScript. The Flu-M1 mutation library was designed with 21 subgroups, each featuring fixed P2 (isoleucine) and P9 (leucine) anchor residues, two additional residues fixed per subgroup, and the remaining positions mutated with NNK degenerate codons to represent all 20 amino acids. The HLA-A*02 fully randomized library was designed by fixing P2 (isoleucine, leucine, or methionine) and P9 (valine, methionine, or leucine) and randomizing all remaining positions using NNK degenerate codons to encode all possible amino acids. These libraries were synthesized as degenerate primers.

Oligonucleotides from the Flu-M1 mutation library were mixed at an equal molar ratio. For other libraries, oligonucleotides were directly subjected to PCR using a reverse primer to generate double-stranded DNA (dsDNA) libraries, which were purified by gel extraction. The dsDNA libraries were assembled with the BsmBI-linearized pHAGE-BsmBI-SCD-IRES-LNGFR vector via Gibson assembly^49^. The assembled vector was purified and electroporated into E. cloni 10G competent cells (Lucigen) at 1,800 V.

#### Genomic DNA extraction, amplification, and NGS sequencing

Genomic DNA (gDNA) from sorted Jurkat cells in PBS was purified using the Magen gDNA Extraction & Purification Kit following the manufacturer’s protocol. For screening without TCR-SCD linking, two rounds of PCR amplification were performed on the extracted gDNA using KOD One polymerase (TOYOBO). After genomic amplification and the addition of indexes and adapters, DNA was pooled and sequenced by Novogene using paired-end 150-bp (PE150) sequencing with at least 1 Gb of clean data. For screening the linked TCR–SCD library, gDNA was extracted and amplified with the corresponding primers, and 10 μg of purified DNA per round was used for library preparation and sequenced by Novogene on a PacBio Revio instrument.

#### SEEKER screen against the Flu-M1 mutation library

For rounds 1 and 2, 2.2×10⁷ J23-A*02 cells expressing Flu-M1-TCR were transduced with the Flu-M1-mut-LV library at an MOI of 20, then seeded in hydrogel for 48 h. ZsGreen⁺ cells were sorted using a BD FACSAria III, and genomic DNA was extracted; SCD sequences were amplified by nested PCR and subcloned into the BsmBI-linearized pHAGE-BsmBI-SCD-IRES-LNGFR vector to construct the next-round plasmid library. For round 3, to minimize multicopy carriers, 5×10⁷ J23-A*02 cells expressing Flu-M1-TCR were transduced with the round-2 subcloned library at an MOI of 0.3; 3.5×10⁵ cells were sorted, and their genomic DNA was pooled for amplification. Amplification products from all rounds were sequenced. For data visualization, the top 200 sequences from round 3, ranked by read count, were filtered to exclude those containing stop codons (n = 3). The remaining 197 sequences were translated and deduplicated at the amino-acid level, yielding 126 unique peptides, which were pooled to generate amino acid usage frequency plots and peptide network plots.

#### SEEKER screen for DMF5-TCR against the HLA-A*02 fully randomized library

For rounds 1 and 2, 2.5×10⁷ J23-A*02 cells expressing DMF5-TCR were transduced with the A*02-NNK library (round 1) or the round-2 subcloned library at an MOI of 20. For round 3, 2.5×10⁷ cells were transduced at an MOI of 0.3. Libraries from all rounds were sequenced. For data visualization, the top 300 sequences from round 3, ranked by read count, were translated and deduplicated at the amino-acid level, yielding 295 unique peptides, which were pooled to generate amino acid usage frequency plots and peptide network plots.

#### SEEKER screen for four TCRs against the HLA-A*02 fully randomized library

J23-A*02 cells were separately transduced with lentiviruses encoding the DMF5, 1G4, EBV, or Flu-M1 TCR, and equal numbers of cells (2.5×10⁷ each) were pooled. For rounds 1 and 2, the pooled cells (1×10⁸ total) were transduced with the A*02-NNK library (round 1) or the round-2 subcloned library at an MOI of 20. For round 3, prior to transduction, 1×10⁸ total cells (2.5×10⁷ cells per TCR) were labeled with CellTrace Violet (CTV) and CellTrace Far Red (CTR) to enable multiplex sorting: DMF5-TCR (CTV⁻CTR⁻, lower left quadrant), 1G4-TCR (CTV⁺CTR⁺, upper right quadrant), Flu-M1 TCR (CTV⁻CTR⁺, lower right quadrant), and EBV TCR (CTV⁺CTR⁻, upper left quadrant). After washing, cells were pooled, transduced at an MOI of 0.3, and activated cells were gated and sorted into four TCR-specific populations. Amplification products from rounds 1–2 and TCR-specific round 3 sorts were sequenced. For data visualization, the top 200 barcodes ranked by round-3 read count for each TCR were filtered to exclude barcode duplication, those containing stop codons and those with R3/R2 fold changes below thresholds. A total of 147, 118, 104, and 30 unique peptides, for DMF5, 1G4, EBV and Flu-M1, respectively, were used to generate amino acid usage frequency plots.

#### T cell activation by peptide-pulsed antigen-presenting cells (APCs)

Single peptides (99% purity) were synthesized by GenScript. K562-A*02 cells were used as APCs for the assay. For 1G4 TCR, 293T-A*02 cells were used as APCs because 1G4 reacts with endogenous antigen of K562 cells. In brief, 3×10⁴ APCs and 3×10⁴ TCR-expressing cells were mixed in a 96-well plate, and peptides at specific concentrations were added. After 48 h, T cells were gated and the percentage of NFAT-zsGreen activation was analyzed by flow cytometry.

#### In-droplet dual-asymmetric overlap-extension PCR (idaOE-PCR) for linking TCR and pMHC

To physically link TCR and SCD sequences carried by the same TAPCell, sorted cells (1×10⁵–1×10⁶ per reaction) were washed, resuspended in 85 μL of 0.25 M sucrose, and heated at 80°C for 40 min for thermal lysis. After cooling, the suspension was mixed with 5 μL of NotI-HF (NEB) and 10 μL of rCutSmart Buffer and incubated at 37°C for 2 h. Digested cells were washed once with 1 mL of 0.25 M sucrose and resuspended in a modified 2 mL KOD emulsion-PCR mix containing 1 mL of 2× KOD One Master Mix (TOYOBO), 32 μL of ET-SSB (NEB), 200 μL of 2 mM dNTPs (TOYOBO), 20 μL of 10% poloxamer 188 (Thermo Fisher), 240 nM outer primer pairs, 18 nM inner overlap primer pairs, and nuclease-free water (Thermo Fisher); EvaGreen was included only in reactions intended for droplet-amplification imaging. Emulsions were generated using a Bio-Rad ddPCR cartridge, loading the equivalent of 800–12,000 cells per 20 μL sample well, transferred to a PCR plate, and subjected to emulsion PCR (95°C for 2 min, followed by 52 cycles of 95°C for 30 s, 60°C for 30 s, and 68°C for 3 min; ramp rate 2°C/s).

Following amplification, 20 μL of TE buffer, 1 μL of Proteinase K, and 3 μL of DNA loading dye (Tiangen) were added to each well to terminate the reaction and label the aqueous phase. Emulsions were broken by adding 80 μL of 40% PFO/HFE-7500 (Sigma-Aldrich) per well and vortexing. The plate was centrifuged at 2,000 ×g for 4 min and incubated at 55°C for 20 min for Proteinase K digestion. Samples were then pooled into 15 mL tubes and centrifuged at 2,000 ×g for 4 min to separate the fluorinated oil from the aqueous phase. The entire aqueous phase was collected and purified using a DNA purification kit (Magen). Purified DNA was resolved by agarose gel electrophoresis, and the ∼1,930-bp band was excised and gel-purified (Magen). The recovered DNA was used either to amplify the TRAV–peptide region for NGS sequencing or as the template for semi-nested PCR of the full-length TRB– TRA–peptide cassette to increase DNA yield.

For first-step library construction, 3 μg of nested-PCR product was assembled with 6 μg of the linearized pHAGE vector carrying a partial SCD tail using NEBuilder HiFi DNA Assembly Master Mix (NEB) in a 600 μL reaction. The assembly product was desalted and electroporated into 12 vials of E. cloni 10G SUPREME electrocompetent cells (Lucigen), and the first-step library plasmid was extracted using a plasmid extraction kit (Magen). To reconstitute the full-length cassette, 60 μg of the first-step library plasmid, in which the high-GC linker is flanked by two MluI sites, was digested with 60 μL of MluI-HF (NEB) and gel-purified to yield the linearized library vector; 9 μg of this vector was then assembled with 1.5 μg of the TRAC-T2A-SCD fragment using NEBuilder HiFi DNA Assembly Master Mix. The assembly product was desalted and electroporated into 12 vials of E. cloni 10G SUPREME cells, and the resulting second-step library plasmid was extracted and served as the plasmid library for subsequent screening rounds. Each cloning step maintained >3×10⁸ transformants to minimize library loss.

#### Single-cell RNA sequencing and analysis of joint samples

Samples were prepared according to the 10x Genomics Chromium protocol^50^ and submitted to Accuramed Technology Ltd (Shanghai) for sequencing. Single-cell RNA-seq data from six samples (AJSS-1, AJSS-2, ASSS-2, AFSS-3, AJSS-3, and AJSS-4) were analyzed using Seurat (v5.3.0)^51^ in R (v4.4.0). Cells were filtered by mitochondrial and hemoglobin gene percentages (<5% each) and minimum detected genes (≥300), and doublets were removed using DoubletFinder^52^ algorithm. Gene counts were log-normalized (scale factor 10,000), the top 3,000 highly variable genes were selected and scaled for principal component analysis (50 PCs), and batch effects across samples were corrected using Harmony (v1.2.3) ^53^with sample identity as the grouping variable. Clustering (Louvain algorithm, resolution 1.0, selected from 0.1– 1.6 based on marker expression and biological interpretability) and UMAP visualization were performed on the Harmony-corrected embedding.

#### T cell isolation and annotation

T cells were extracted from the quality-controlled dataset based on Human Primary Cell Atlas (HPCA) annotation^54^ and manual curation; cells annotated as NK cells were removed, and clusters with high expression of HBB, myeloid markers (S100A8/S100A9, LYZ), or the mast cell marker KIT were iteratively excluded, yielding a final dataset of 18,415 T cells. Subtypes were assigned based on canonical markers identified using FindAllMarkers (Wilcoxon rank-sum test; only.pos = TRUE, min.pct = 0.25, logfc.threshold = 0.25), including lineage (CD3D, CD4, CD8A), differentiation (CCR7, TCF7), and effector (GZMB, GZMK, NKG7) markers and lineage-defining transcription factors (FOXP3, EOMES), covering CD8⁺ and CD4⁺ naive and memory subsets, Treg, MAIT, γδ T (including cytotoxic), and T cell stress response (TSTR) cells. The remaining S100A8/A9⁺ cluster was removed after confirmation of myeloid contamination.

#### TCR clonotype analysis

TCR sequences were recovered from matched single-cell V(D)J libraries and assembled into full-length TCRα/TCRβ sequences. Clonotypes were defined by paired CDR3 amino acid sequences (CTaa), with single-chain clonotypes merged when the matching chain was identified; clone size was categorized as non-expanded (1 cell), small (2-5), medium (6-20), large (21-100), or hyperexpanded (>100) cells and visualized across subtypes on UMAP embeddings. Figures were generated using ggplot2 (v3.5.2) and assembled with patchwork (v1.3.1).

#### TCR and HLA-B*27-restricted epitope library construction

Of note, TCR005 and TCR101 were from a single T cell that harbor two productive TRA rearrangements paired with a single TCRβ chain. Because both α chains may form functional receptors with the β chain, we treated the two pairs as separate TCRs in the screen.

The sequences of 101 highly expanded TCRs from AS samples were extracted and reconstructed into full-length single-chain TCRβ-furin-SGSG-P2A-TCRα constructs. Individual plasmids harboring these TCR sequences were synthesized by GenScript. TCR plasmids within each HLA allele group were pooled at an equal molar ratio, and balanced representation of each TCR was verified by NGS. The pooled plasmids were electroporated into competent cells, and the TCR library plasmids were extracted to prepare the TCR-expressing lentiviral library.

For the HLA-B*27 epitope library, human proteome sequences and annotated viral sequences (TaxID: 10239) were downloaded from UniProt^55^. HLA-B*27 epitopes from both the human and viral proteomes were predicted using NetMHCpan-4.1^56^ with default parameters. In total, 1,152,013 human self-peptides and 432,245 virus peptides were synthesized as oligonucleotides with flanking sequences (GenScript). After extracting plasmids from the cloned HLA-B*27 library, 100 ng of plasmids were amplified using sequencing primers to validate cloning efficiency, with >99.9% of library candidates recovered from sequencing data.

#### HLA-B*27-restricted library screening

Two screening experiments were performed (for B*27:04 and B*27:05), both based on the idaOE-PCR linked-library workflow described above. J23-B*27 cells were first transduced with a pooled 50-TCR library. Transduced cells were sorted for TCR expression, and genomic DNA was extracted from 1×10⁵ cells for sequencing to verify the uniformity of TCRA and TCRB expression.

Subsequently, after expansion, 6×10⁸ HLA⁺ cells expressing 50 TCRs were transduced with concentrated lentivirus at an MOI of 20 in the presence of 400 nM dasatinib. After washing, cells were cultured in hydrogel for 48 h and sorted for

ZSGREEN⁺ cells. Sorted cells were subjected to the idaOE-PCR and cloning procedures described above to obtain the library for subsequent screening rounds.

For round 2 and subsequent rounds, 2×10⁸ HLA⁺ cells were transduced with concentrated lentivirus at an MOI of 1. Cells were cultured in hydrogel for 72 h and sorted for ZsGreen⁺ cells; genomic DNA was extracted from sorted cells, and full-length TCR–SCD cassettes were amplified and cloned into the pHAGE vector to obtain the next-round library. Amplicons from rounds 2 and 3 were subjected to PacBio full-length TCR–SCD sequencing.

#### Validation of screening results in the SCD format

For each library-on-library screen, the top hits were first selected and further refined by incorporating peptide-sharing motif similarity. Peptide DNA coding sequences were synthesized as complementary oligonucleotides with sticky ends, annealed, ligated into the pHAGE-SCD vector, and transformed to generate single-antigen SCD-form plasmids. Individual lentiviruses were packaged in 96-well plates, and 50 μL of each lentivirus was used to transduce 2.5×10⁴ HLA-expressing cells carrying the corresponding TCR. After 48 h, PE-LNGFR staining was used to confirm viral infection efficiency, and flow cytometry analysis was performed to determine cell activation rates.

#### Analysis of viral–self peptide mimicry for TCR59 and TCR63

Viral and self peptides identified in the screen were compared in all virus–self combinations with a peptide-length difference ≤ 6 residues. Sequence similarity was quantified by a combined score of 0.6 ×Levenshtein similarity + 0.4 ×3-mer Jaccard index, where Levenshtein similarity = 1 − (edit distance / length of the longer peptide) and the Jaccard index was computed on 3-mer sets of the two peptides. A pair was called when the combined score exceeded 0.35 or when the two peptides shared a contiguous motif of ≥ 4 amino acids. This yielded 437 mimicry pairs, of which 60 involved TCR59 virus peptides and 377 involved TCR63 virus peptides.

### QUANTIFICATION AND STATISTICAL ANALYSIS

#### Raw sequence processing and read counting

Illumina NGS data were processed with custom Python scripts on a Linux server: regular expressions were used to filter valid sequences, and read counts were calculated for each round of screening. PacBio sequencing data were first converted from BAM to FASTQ format using SAMtools^57^. A custom Python workflow was used to parse TCR and peptide sequences, map peptides to the designed library, and identify point mutations through minimap2-based alignment^58^.

Sequences carrying frameshift mutations were discarded, while sequences matching the designed library (allowing isolated synonymous mutations) were retained.

Relative percentage of each peptide hit was calculated by dividing individual read counts by the total count. Results were visualized using Python plotting libraries.

#### Statistical analysis

Statistical analysis was performed using Python and Excel. For endpoint comparisons, two-tailed Student’s *t* tests were used.

## Acknowledgments

The work was funded in part by the Fundamental and Interdisciplinary Disciplines Breakthrough Plan of the Ministry of Education of China (JYB2025XDXM603 to H.Q.), National Natural Science Foundation of China (grant T2495270 to H.Q. and X.W. and 32130037 to H.Q.), Tsinghua-Peking Center for Life Sciences, the Beijing Frontier Research Center for Biological Structure and the SXMU-Tsinghua Collaborative Innovation Center for Frontier Medicine. H.Q. is a New Cornerstone Investigator. H.X. is supported by the National Natural Science Foundation of China (Grant No. 82320108010 and 31821003), the National Key R&D Program of China (Grant No. 2018AAA0100302), and the Shanghai Municipal Key Clinical Specialty (Grant No. shslczdzk02602).

## Author contribution

H.Q. conceptualized and supervised the project with clinical input from H.X and data analysis input from X.W. S.L. designed the SEEKER system and conducted the experiments together with L.L., W.W., B-W.L., H.J., Z.G., Y.L., J.W., B.L., L.Z. H.X. supervised patient study and handling of clinical samples. H.Q. and S.L. wrote the paper with input from all authors.

---

**Figure S1.**
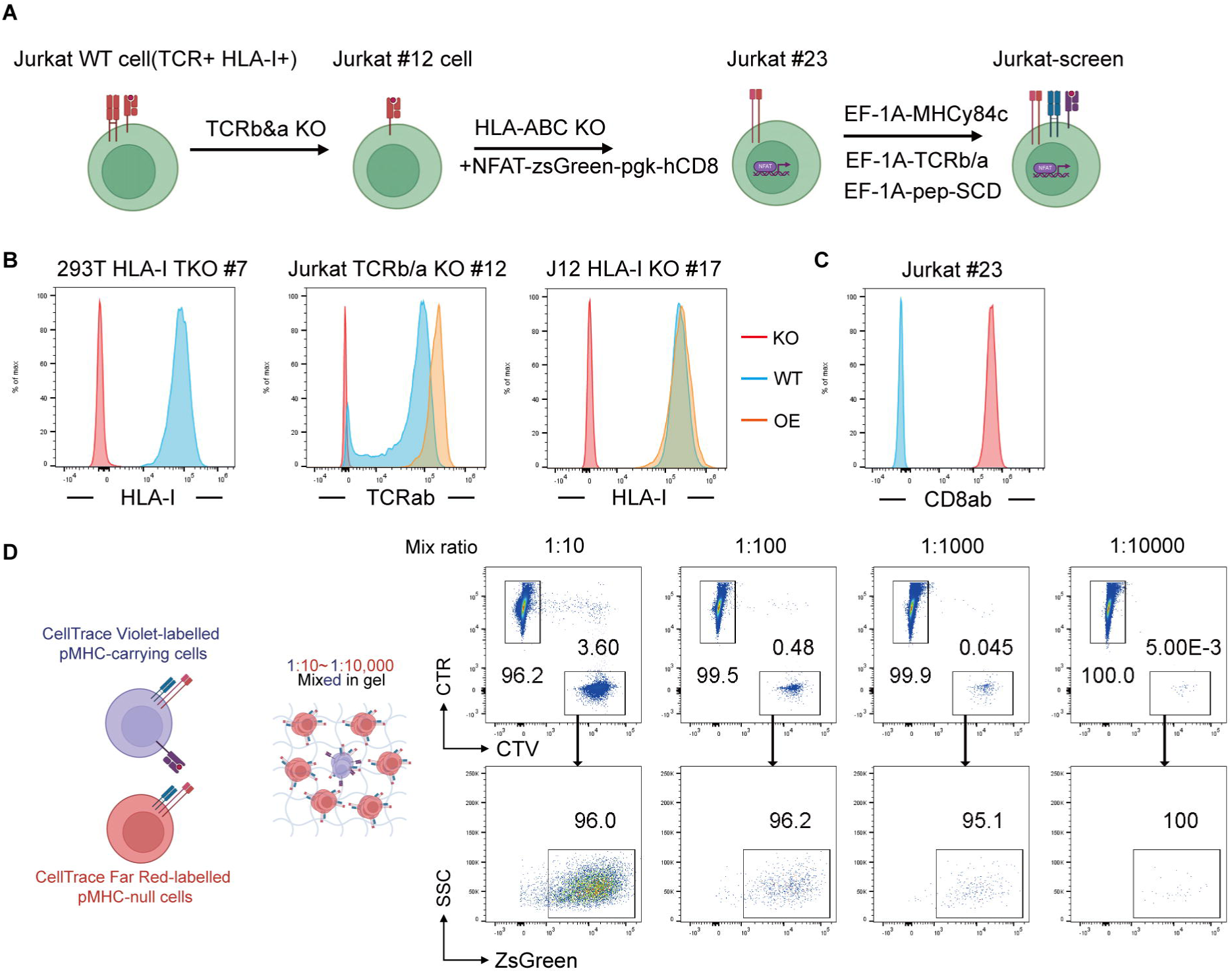
Generation and functional validation of the engineered Jurkat master cell line. (A) Stepwise generation of the Jurkat screen cell line. Wild-type Jurkat cells are first engineered by CRISPR/Cas9 disruption of the endogenous TCRα and TCRβ chains, followed by disruption of the HLA-A, -B, and -C loci and stable integration of an NFAT-ZsGreen reporter with PGK-driven hCD8, yielding the Jurkat master clone; transduction with EF-1α-driven MHC, TCR, and peptide-SCD constructs reconstitutes the screen-competent cell. (B) Flow-cytometry validation of the knockout steps. HLA-I and TCRαβ staining of the edited clones (KO) is overlaid with parental wild-type (WT) and overexpression (OE) controls. (C) hCD8αβ staining of the Jurkat master clone, confirming expression of the introduced co-receptor. (D) Titration of pMHC-carrying cells in hydrogel culture. CellTrace Violet-labeled pMHC-carrying cells are serially diluted from 1:10 to 1:10,000 into CellTrace Far-Red-labeled pMHC-null cells; dye-based gating quantifies the pMHC-carrying fraction, and NFAT-ZsGreen activation is scored across dilutions.

**Figure S2.**
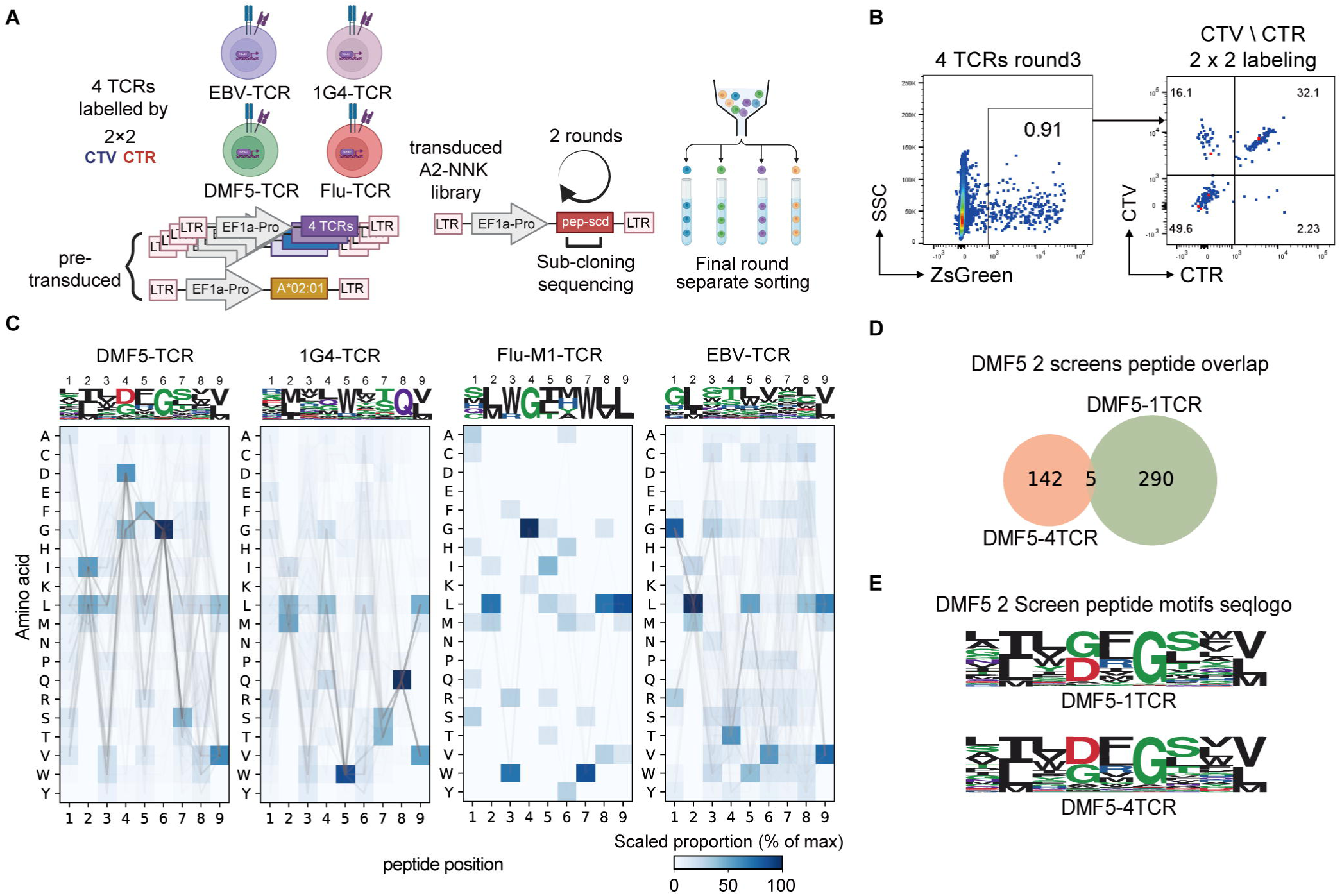
Parallel fingerprinting of multiple TCRs. (A) Schematic of the parallel multi-TCR SEEKER workflow. Multiple TCRs and the MHC heavy chain are pre-introduced into Jurkat master cells before introduction of peptide library. In the final round, activated cells are sorted by dye labeling, followed by sub-cloning and sequencing of peptides only. (B) Final-round flow cytometry profile of the 4-TCR parallel screen. The four TCRs are distinguished by 2×2 CellTrace Violet/CellTrace Far Red labeling. (C) Sequence logos and amino acid usage matrices of the DMF5 (n = 147), 1G4 (n = 118), Flu-M1 (n = 30), and EBV (n = 104) TCRs from the 4-TCR parallel screen. (D) Overlap of the peptide coding DNA sequence sets recovered from the DMF5 single-TCR screen and the DMF5 arm of the 4-TCR parallel screen (Venn diagram). (E) Recognition motifs of DMF5 recovered from the single-TCR screen and the 4-TCR parallel screen, compared as sequence logos.

**Figure S3.**
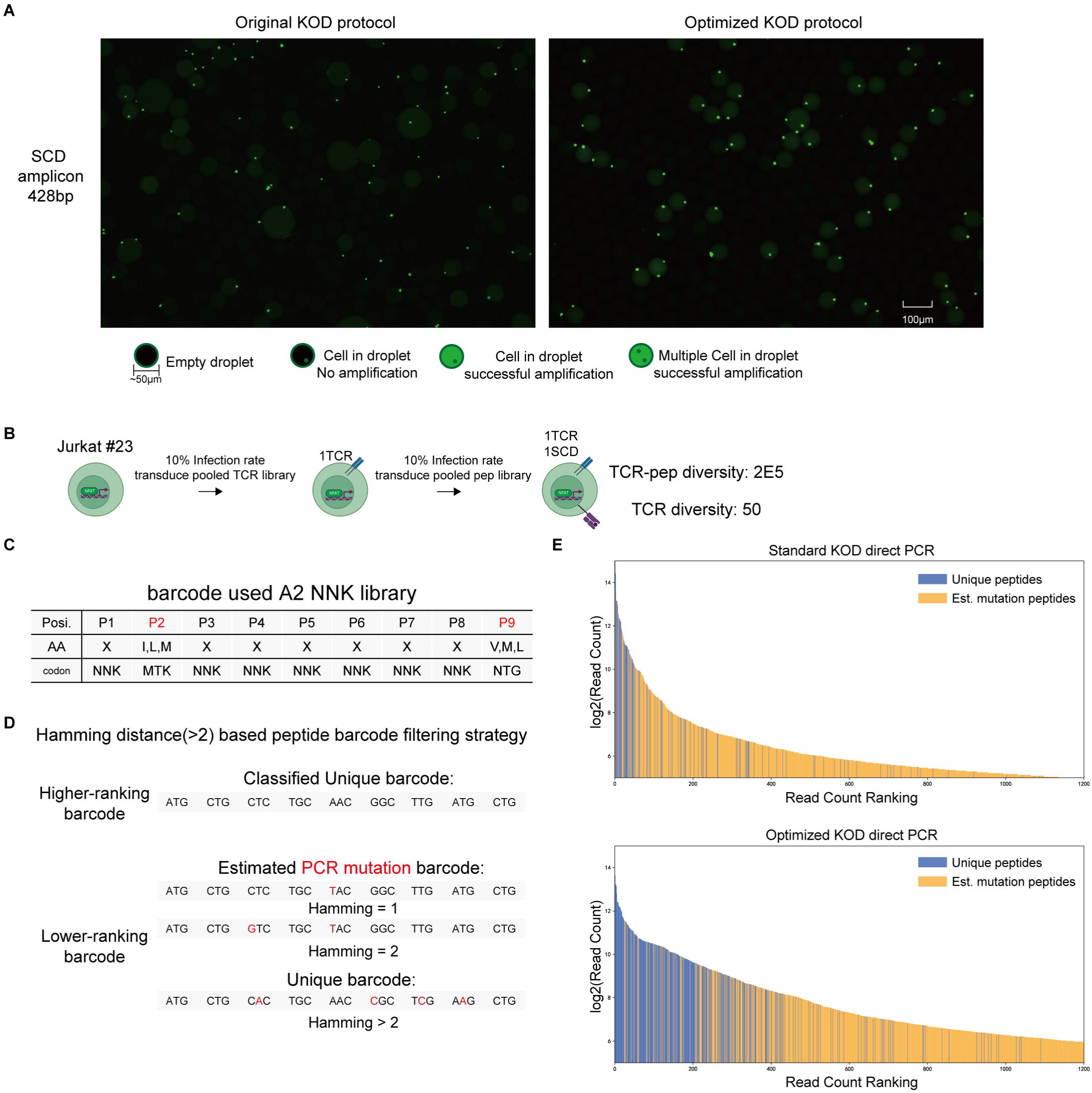
Optimization of idaOE-PCR and quality control of recovery efficiency test. (A) Droplet fluorescence microscopy comparing the original and optimized KOD direct-PCR protocols for amplifying the ∼428bp SCD amplicon. (B) Schematic of the single-copy cell library construction. Jurkat #23 cells are sequentially transduced at a low infection rate with a pooled TCR library and then a pooled peptide-SCD library. (C–D) Scheme of the barcode filtering rules for the idaOE-PCR recovery efficiency test. (C) The HLA-A*02-restricted NNK peptide library was used as sample barcodes. (D) Barcodes (27 nt) were ranked by read count, and each was compared nucleotide-by-nucleotide to all higher-ranking unique barcodes; those within a Hamming distance of ≤2 nt from a higher-ranking barcode were considered PCR-introduced variants, whereas those differing by >2 nt from all established ones were counted as unique. (E) Read-count rank distributions of the resulting unique peptides and filtered variants under the standard and optimized KOD direct-PCR conditions.

**Figure S4.**
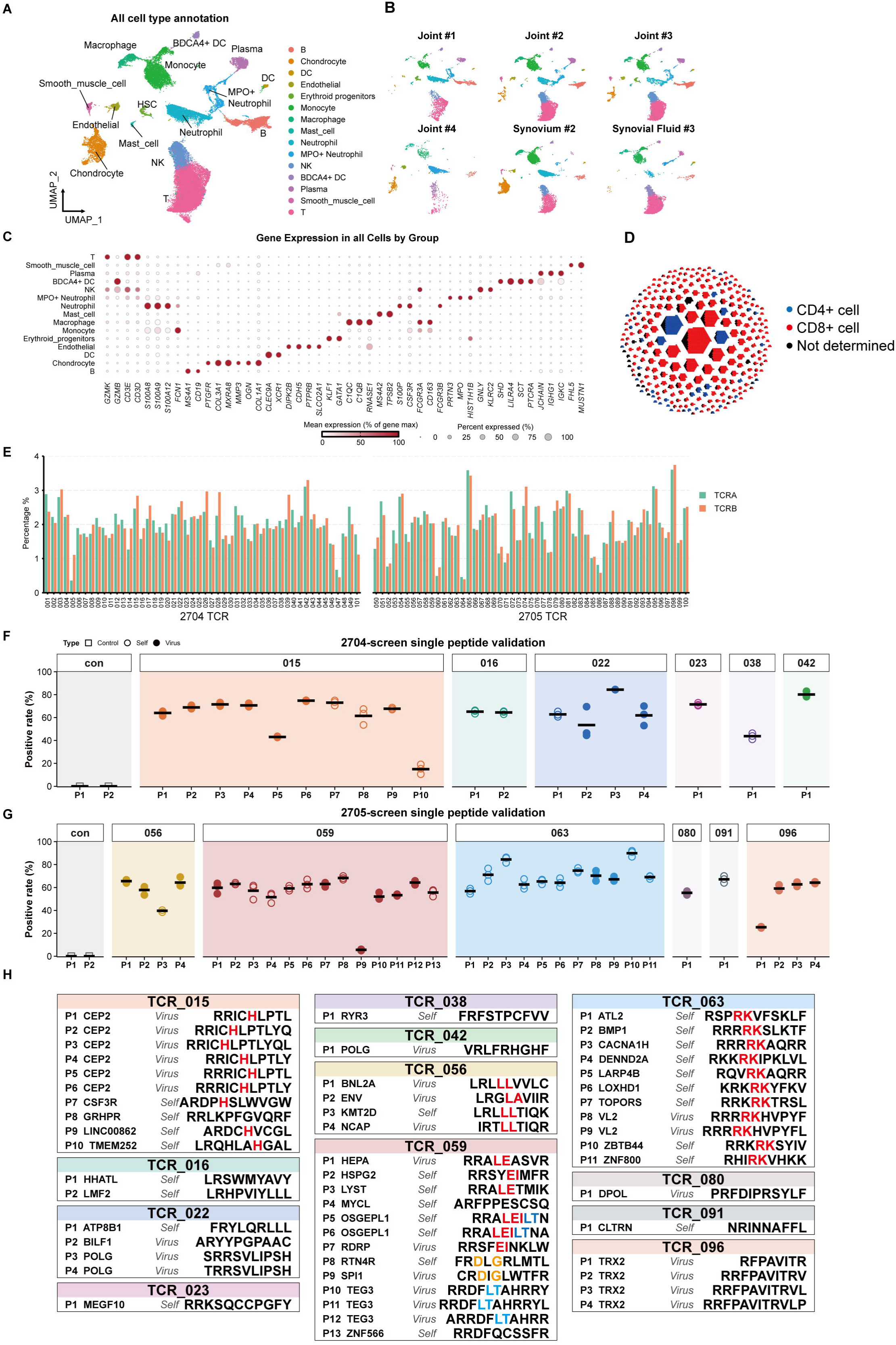
Single-cell atlas of AS joint tissues and validation of screened TCR– peptide pairs. (A) UMAP visualization of the combined joint-tissue atlas with all cell populations annotated. (B) UMAP visualization of the atlas split by tissue and patient sample. Sampling sites: Joint (AJSS), Synovium (ASSS), Synovial Fluid (AFSS). (C) Canonical marker-gene expression across all annotated cell populations. (D) Honeycomb plot of expanded TCR clonotypes by CD4/CD8 phenotype. (E) Representation of individual TCRs in the pooled AS-TCR expressing cell library. The percentages of cells expressing the TCRα and TCRβ chains of each TCR clone are shown for the HLA-B*27:04 and HLA-B*27:05 groups, confirming low-MOI transduction and balanced library expression. (F and G) Single-peptide validation of candidate TCR–peptide pairs from the HLA-B27:04 screen (F) and the HLA-B27:05 screen (G). Each reactive TCR was individually tested against its self-derived and virus-derived candidate peptides, with J23-B*27-50 TCR cells stimulated by the irrelevant peptide SCD serving as a negative control. Bars indicate the means of three replicates per condition. (H) Summary of the peptides validated for each reactive TCR, listing peptide identifiers, source genes, self-or virus-derived origin, and sequences; sequence similarity between the cross-recognized viral and self epitopes is labelled by colors.

**Figure S5.**
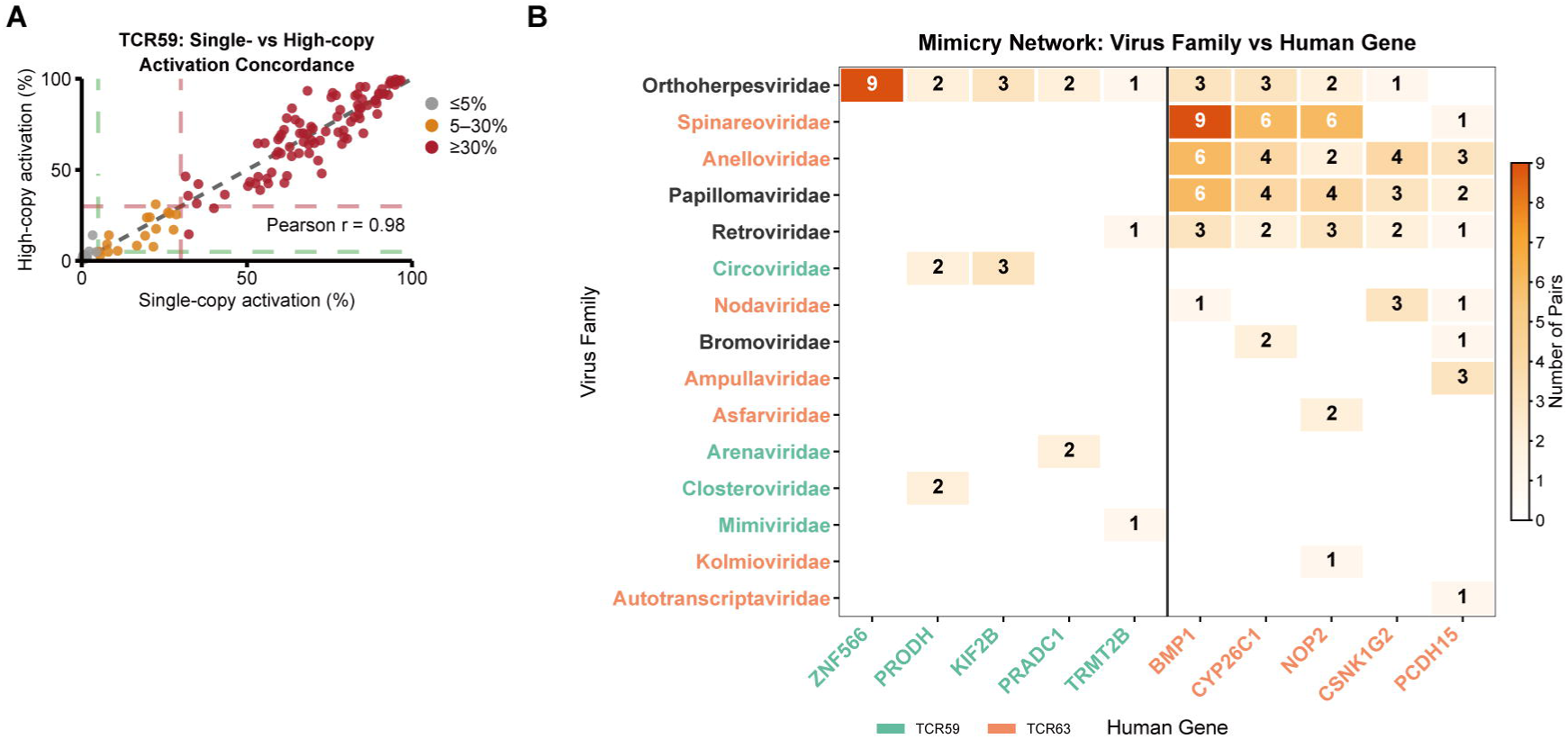
Concordance across TCR expression formats and summary of virus– self mimicry pairs. (A) Concordance of TCR59 peptide (n=192) validation between single-copy and high-copy TCR expression formats. Each point represents one peptide; point color indicates the single-copy activation class, and dashed lines mark the activation thresholds. (B) Mimicry-participating virus–self peptide pairs summarized by viral family and human gene for TCR59 and TCR63. Numbers indicate pair counts. For each TCR, the five self-antigen genes with the most pairs are shown. Pairs were defined as described in the Methods. Only peptides activating their TCR (> 5%) were compared.

